# Variability of axon initial segment geometry and its impact on hippocampal pyramidal cell function

**DOI:** 10.1101/2024.12.16.628625

**Authors:** Nikolas Andreas Stevens, Maximilian Achilles, Juri Monath, Maren Engelhardt, Martin Both, Christian Thome

## Abstract

Action potentials, the primary information units of the nervous system, are usually generated at the axon initial segment (AIS). Changes in the length and position of the AIS are associated with alterations in neuronal excitability but there is only limited information about the baseline structural variability of the AIS. This work provides a comprehensive atlas of the diversity of proximal cell geometries across all anatomical axes of the murine hippocampus, encompassing dorsal-ventral, superficial-deep, and proximal-distal regions. We analyzed the morphology of 3,936 hippocampal pyramidal neurons in 12 animals of both sexes, focusing on AIS length, position, and their association with proximal cellular features such as the soma and dendritic geometries. Notably, neurons with axon-carrying dendrites were significantly more common in ventral compared to dorsal hippocampal areas, suggesting a functional adaptation to regional demands. Validation of this finding in human samples confirms the translational relevance of our murine model. We employed NEURON simulations to assess the functional implications of this variability. Here, variation in proximal geometry only minimally contributed to neuronal homeostasis, but instead increased heterogeneity of response patterns across neurons.

## INTRODUCTION

Neuronal populations are often separated and categorized by common morphological characteristics, such as the geometry of the soma, axonal arborization or the shape of the dendritic tree. While practical, this approach reinforces a simplified concept of a “typical neuron”, disregarding the substantial diversity within individual regions and subregions. In fact, the “typical neuron” may constitute an exception rather than the rule in a complex and varied neuronal landscape.

The shape of the cell body and nearby structures has profound functional implications, particularly in how neurons integrate inputs and generate firing patterns (Gouwens et al., 2020, Jiang et al., 2015, Markram et al., 2015, Peng et al., 2021). The axon initial segment (AIS) plays a pivotal role in this context. Due to its high expression of voltage gated ion channels (Hodgkin and Huxley, 1952, Kole and Stuart, 2012, Leterrier, 2018) and preferable electrotonic conditions, most action potentials are generated at the AIS (Baranauskas et al., 2013, Foust et al., 2010, Hu et al., 2009b, Kim et al., 2005, Kole and Stuart, 2012, Meeks and Mennerick, 2007, Palmer and Stuart, 2006, Royeck et al., 2008, Shu et al., 2007). Remodeling of the AIS, as well as mutations in its scaffolding proteins, have been associated with mental pathologies such as bipolar disorder, epilepsy, intellectual disability, autism spectrum disorder, and stroke (Fujitani et al., 2021, Huang and Rasband, 2018, Stoler and Fleidervish, 2016). However, several studies have demonstrated that the length of the AIS and its distance from the soma undergo plastic changes in response to alterations in neuronal activity. This AIS plasticity in turn correlates with functional changes in the respective cells (Grubb and Burrone, 2010, Jamann et al., 2021, Kuba, 2012). Computational studies suggest that the AIS geometry directly impacts neuronal excitability, firing precision, and network behavior (Goethals and Brette, 2020, Gulledge and Bravo, 2016, Hamada et al., 2016, Kole and Brette, 2018), but see also (Moubarak et al., 2019). However, it remains unclear how AIS plasticity relates to the natural variability of initial segments. Longer AIS are associated with elevated neuronal excitability due to their increased area for voltagegated ion channels, which are critical for triggering action potentials. In the avian auditory system, AIS lengths and positions vary according to sound frequency (Kuba, 2012). The impact of AIS location within the axonal arbor on excitability is multifaceted; more proximal locations are reached faster by somatic depolarization. On the other hand, the substantial capacitive sink provided by the large somatic membrane is thought to reduce neuronal excitability, rendering distal AIS locations again more excitable than proximal ones (Goethals and Brette, 2020, Gulledge and Bravo, 2016, Kole and Brette, 2018).

The situation becomes even more complex when the AIS emerges from a dendritic branch, creating an electrically privileged site for synaptic input (Hamada et al., 2016, Hofflin et al., 2017, Thome et al., 2014). Inputs received at these axon-carrying dendrites (AcD) have privileged access to the AIS, especially during network states in which perisomatic inhibition limits the efficiency of regular excitatory input. Thus, most CA1 pyramidal neurons that fire during sharp wave ripple oscillation have dendritic axon origins (Hodapp et al., 2022). Neurons with AcD morphology (AcD cells) in ventral CA1 also receive also more synaptic innervation from the contralateral hemisphere (Stevens et al., 2024). In hippocampal cultures, AcD cells showed less AIS plasticity compared to neighboring nonAcDs in response to induced network changes (Han et al., 2025). This illustrates that divergent AIS geometries correlate with functional roles of individual neurons within the same cell population. Electrophysiological recordings have demonstrated that neurons with AcD morphology are indistinguishable from their counterparts in terms of their basic electrophysiological properties when recorded at the soma (Stevens et al., 2024, Thome et al., 2014). In cortical AcD cells, the diameter of the apical dendrite decreases with AIS distance, thereby normalizing its impact on the action potential threshold and waveform (Hamada et al., 2016). It is thus conceivable that variations in AIS length and positions are counteracted by other morphological parameters such as the size of the soma or apical dendrite (Gulledge and Bravo, 2016).

Occurrence and function of AcD cells were previously studied in well-defined subpopulations, like murine CA1 pyramidal cells of the ventral hippocampus (Hodapp et al., 2022, Stevens et al., 2024, Thome et al., 2014) or cortical layer 5 pyramidal neurons of rats (Hamada et al., 2016). A complete assessment of AcD cell numbers throughout the central nervous system is still pending. However, drawing conclusions even from functionally and structurally related cell populations can be challenging, as these populations often exhibit significant specializations along the anatomical axes. The rodent hippocampus is segregated anatomically, genetically, and functionally into the ventral, intermediate, and dorsal portions along its longitudinal axis (Bannerman et al., 2004, Fanselow and Dong, 2010, Strange et al., 2014). The dorsal and medial sections of the hippocampus are preferentially involved in spatial learning (O’Keefe and Dostrovsky, 1971, Moser and Moser, 1998) and the ventral part plays an important role in mediating anxiety-like behavior and emotional learning (Moser and Moser, 1998, Richmond et al., 1999, Kjelstrup et al., 2002). Place fields are more discrete and precise in the dorsal end of the hippocampus, gradually becoming larger towards the ventral end (Jung et al., 1994, Kjelstrup et al., 2008, Royer et al., 2010). Alongside the behavioral specialization, the anatomical connectivity of these regions is also different. The dorsal hippocampus receives visuospatial information via the medial entorhinal cortex, whereas the ventral hippocampus is highly connected to the amygdala, hypothalamus, and sensory areas (Amaral and Witter, 1989, Fanselow and Dong, 2010, Cenquizca and Swanson, 2007). The intermediate hippocampus receives both, visuospatial input and is also connected to prefrontal and subcortical areas including the amygdala and hypothalamus (Fanselow and Dong, 2010). This overlapping connectivity may allow this region to play a specific role in rapid place learning tasks (Bast et al., 2009). Disease-related activities also differ between both ends. Epileptic seizures were found to originate mainly from the ventral and not the dorsal hippocampus (Akaike et al., 2001, Racine et al., 1977, Toyoda et al., 2013). In contrast, dorsal CA1 pyramidal cells are more prone to damage by ischemia (Ashton et al., 1989). Gene expression studies in rodents also provided evidence for distinct molecular boundaries within the dorsal-ventral axis of the hippocampus (Fanselow and Dong, 2010, O’Reilly et al., 2015). In vitro studies using hippocampal slices showed that CA1 pyramidal neurons gradually increase in excitability from the dorsal to the ventral end of the hippocampus (Milior et al., 2016, Dougherty et al., 2012, Malik et al., 2016). This enhanced excitability towards ventral CA1 encompasses rising firing frequencies, input resistances, action potential thresholds, and resting membrane potentials. At the same time, CA1 neurons in the ventral part have fewer dendritic branches and less dendritic surface area compared to dorsal CA1 neurons. These electrophysiological differences are further accompanied by differential expression of voltage-gated ion channels, such as HCN channels (Dougherty et al., 2013, Marcelin et al., 2012), which play a crucial role in maintaining the resting properties of CA1 neurons and significantly affect the integration and transfer of voltage signals along the somatodendritic axis (Vaidya and Johnston, 2013).

The present study aims to investigate whether AIS morphologies correlate with the established functional specializations along the primary anatomical axes of the hippocampus: transverse (proximo-distal), radial (deep-superficial), and longitudinal (dorsal-ventral). Given the well-characterized differences between these locations in terms of behavioral roles, anatomical connectivity, electrophysiology, and gene expression, it is plausible that populations of pyramidal neurons may exhibit distinct AIS morphologies tailored to the computational demands of their regions. However, no prior research has compared the proximal morphology of neurons across all these axes systematically, nor explored the potential physiological implications of such variations.

## RESULTS

We assessed the proximal cell geometry of hippocampal neurons using the intrinsic fluorescence signal of the commonly used Thy1-GFP mouse line, in which a sparse subset of mature projection neurons are labelled with GFP (Morris and Grosveld, 1989). The location and length of the AIS were visualized using immunofluorescence against the scaffolding protein βIV-spectrin. Regions of interest were defined along the main anatomical axes of the hippocampal formation: **A:** the dorsal, medial, and ventral hippocampal sections (**Figure 1A**), **B:** the hippocampal cell band, divided into the hilus, CA3c, CA3b, CA3a, proximal CA1 (CA1p), central CA1 (CA1m), distal CA1 (CA1d), and subiculum (**Figure 1B**), **C:** along the superficial to deep axis (**Figure 1C**; 20 cells per region per hemisphere, 12 animals, 3-4 months old, both sexes, 3936 neurons in total). Of note, the Thy1-GFP line used in this study (M-line) shows no GFP signal in CA2 so only AIS length measurements were taken here. Since the hilar Thy1-positive cell population is mostly comprised of mossy cells that are morphologically and functionally distinct from pyramidal neurons, we provide them as a resource in our published dataset, but have omitted them from our statistical analysis.

**Figure 1:**
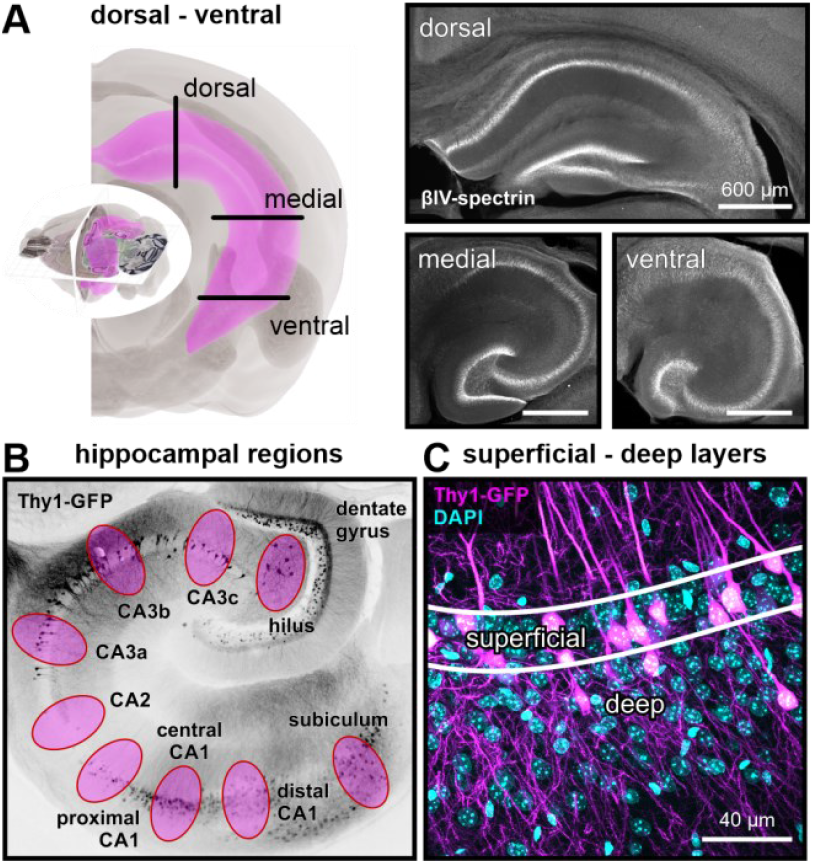
Anatomical axes in the mouse hippocampus. **(A)** The left panel was modified from Allen Brain Explorer and shows the longitudinal orientation of the hippocampal formation in the murine brain. The right panel shows confocal images of AIS labeling along the pyramidal cell band (βIV-spectrin signal) in brain slices obtained in the orientation visualized on the left. **(B)** Confocal image of Thy1-GFP signal in a ventral hippocampal slice. Hippocampal subregions used for analysis are marked in magenta. **(C)** Confocal image of Thy1-GFP and DAPI signals illustrating the separation of the dense ∼2 cell bodies wide superficial layer from the deep layer.

We measured the size of the soma, the diameter of the proximal portions of the apical dendrite, the length, and position of the AIS, and in the case of dendritic axon onsets, the length and diameter of the AcD stem dendrite separating soma and axon-dendrite bifurcation (see fluorescent images in **Figure 2**). Our experimental outline was not designed to study the impact of sex or brain hemisphere, but when we compared the median values for each parameter in each hippocampal location, we found no significant differences between those groups. Since we focused our study on intrinsic variability and interactions, we collected an equal number of geometries across all anatomical subregions (20 cells) and pooled them for further analysis.

**Figure 2:**
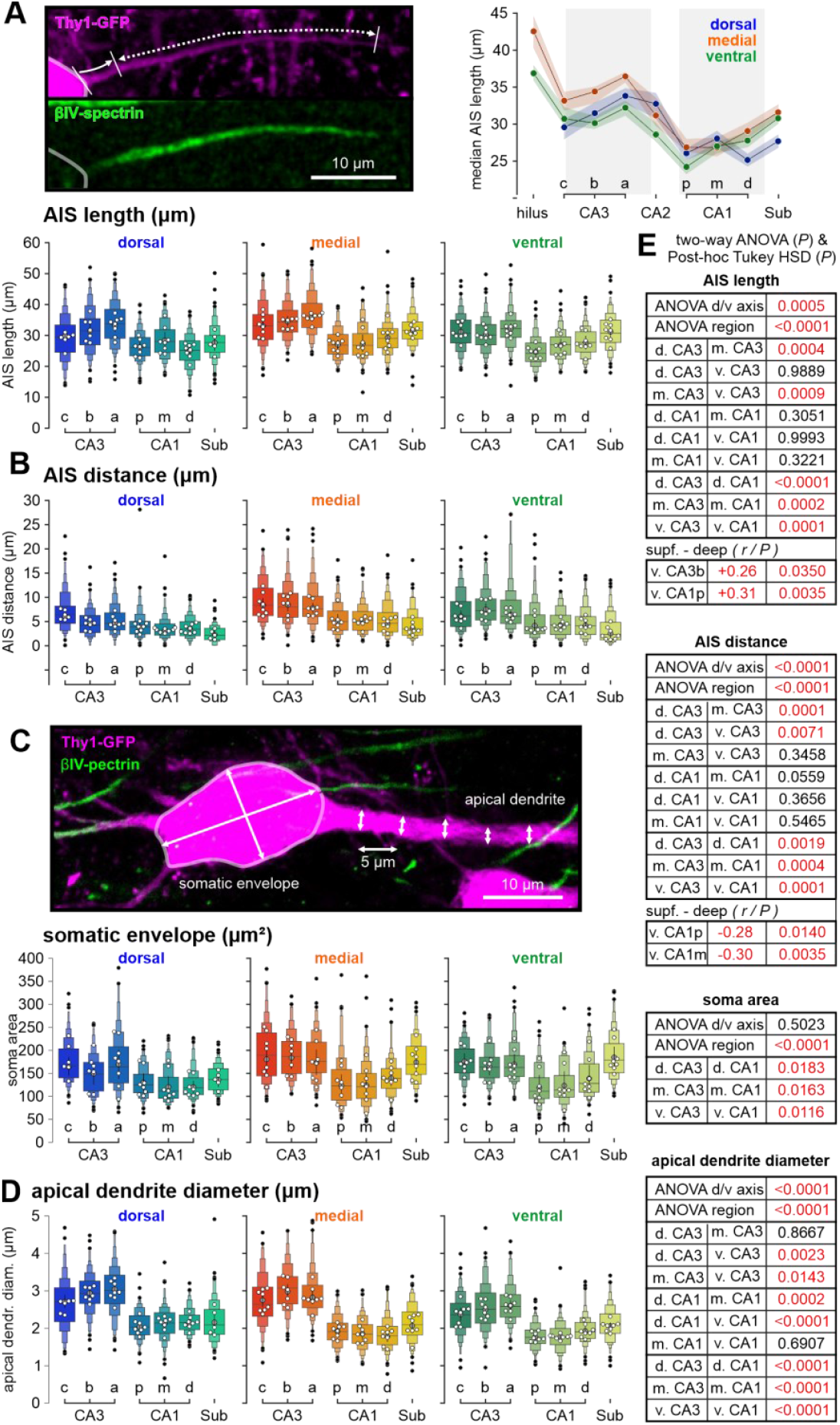
Proximal cell geometry in hippocampal subregions. **(A)** Confocal image of intrinsic GFP label (magenta) and immunolabeled AIS (βIV-spectrin, green). Arrows show length and distance of βIV-spectrin signal within the axon. The right panel shows the length distribution of AIS across the hippocampal cell band of the dorsal (blue), medial (orange), and ventral (green) hippocampal formation. Bottom panel: extended box plots visualize length distributions of the AIS across the different subregions of the hippocampal formation. Median values given in white, mean of medians with SD in grey, and computed outliers in black (see methods for details on letter-value plots, some outliers, <10 in whole Figure 2 & 3, are cut for clarity). **(B)** Plots visualize the distribution of AIS distances from the soma across different subregions of the hippocampal formation. **(C)** Confocal image of intrinsic GFP label (magenta) and immunolabeled AIS (βIV-spectrin, green). Soma size was captured as an approximated ellipsoid by measuring the longitudinal length and diameter of the soma. Bottom plots visualize soma area distributions across the different subregions of the hippocampal formation. **(D)** Extended box plots visualize distributions of apical dendrite diameters across different subregions of the hippocampal formation. Values are the means from 5 consecutive width measurements with 5 µm spacing (see confocal image in C). **(E)** Statistical comparison between hippocampal areas and sections across the dorsal to ventral axes of the hippocampus. P values are derived from two-way ANOVA tests, following Tukey post-hoc analysis (n=hippocampi). Correlations across the superficial to deep axes were analyzed using Spearman correlation followed by Bonferroni correction across all subregions (n=cells). Note, the majority of weak correlations is suppressed by this approach. Dataset derived from 3936 cells from 12 animals in all figures)

### AIS length & distance

Several studies demonstrated plastic changes in AIS length and distance to the soma in response to changes in neuronal activity. We mapped the length of AIS throughout the hippocampus using the marker βIV-spectrin **(Figure 2A, Figure S1)**. Initial segments were largest in the hilus region. Along the pyramidal cell band, median AIS lengths increased along the CA3 axis until they reached the highest values in CA3a. Their length then drops to the lowest at proximal CA1, with CA2 pyramidal cells showing intermediate levels. Across the dorsal to ventral axis, the AIS were longest in the intermediate portion of the hippocampus. Along the superficial to deep axis, AIS were longer in the deeper layers, most prominently in dorsal CA3b and the proximal portion of medial CA1 (*r*-value: +0.26 and +0.31, respectively).

AIS distance was defined as the spatial separation of the soma and the start of the βIV-spectrin signal **(Figure 2A, Figure S1)**. Dendritic AIS origins were included in this analysis (compare **Figure S2**). The initial segment label often started at a considerable distance from the soma across all hippocampal regions. AIS distances generally decreased from CA3 towards the subiculum in all hippocampal planes. The most significant reduction was again seen from CA3a to CA1p, where AIS onsets were about 2 µm shorter. AIS distances were largest in the medial hippocampus and shortest in the dorsal planes. Across the superficial to deep axis, the AIS onset moved closer towards the cell body, most prominently in ventral CA1 in the middle and proximal portions (*r*-value: −0.30 and −0.28, respectively).

Computational studies suggest that the distal end of the AIS compartment is a better predictor of neuronal excitability than its proximal start or total length (Goethals and Brette, 2020). Therefore, we compared the distal end of the AIS between hippocampal pyramidal cells across all anatomical axes (**Figure S3**). We found that the distribution of AIS endpoints (AIS distance + AIS length) follows the pattern observed for AIS lengths very closely. AIS length and distance correlated negatively in CA3 and subicular neurons of medial and ventral planes (medial CA3: *P* < 0.001, medial subiculum: *P* = 0.0035, ventral CA3: *P* = 0.0005, ventral Sub: *P* = 0.0818, pairwise Spearman correlation). This demonstrates that distal AIS location may be partially compensated by shorter AIS.

### Soma and apical dendrite

The somatic membrane provides a considerable sink for synaptic currents which makes cell size a strong predictor of neuronal excitability in general. However, it also interacts directly with spike initiation at the AIS through electric interactions such as resistive coupling (Goethals and Brette, 2020, Gulledge and Bravo, 2016). We found somata of neurons in the dorsal hippocampal areas to be smaller compared to the medial and ventral hippocampus (**Figure 2C**). However, the largest differences occurred between hippocampal areas within each cutting plane. Somata of CA3 neurons are larger than CA1 cells, while subicular neurons show mid-range levels in dorsal and CA3-like levels in medial and ventral planes. The diameter of the proximal apical dendrite was measured as the mean of five consecutive diameters with 5 µm spacing (**Figure 2D**). Their distribution mirrors the pattern found for soma size. Apical dendrites in the dorsal hippocampus were thinner compared to their medial and ventral counterparts. The main hippocampal areas showed the strongest regional differences, with the largest apical dendrite diameters found in CA3 cells.

### Dendritic axon origins

Cells with dendritic axon origins are very common in the ventral CA1 of the mouse hippocampus, comprising around half of all pyramidal cells (Thome et al, 2014; Stevens et al., 2023). However, in vivo investigations targeting cells in the dorsal CA1 found much lower levels of this morphology (Hodapp et al., 2022). When we quantified the propensity for this morphology across the whole hippocampus, we found that AcD cells are common across all hippocampal regions (**Figure 3A & 3B**), but with significant and large differences in distribution across all anatomical axes. The dorsal hippocampus featured generally less dendritic axon origins compared to medial and ventral planes. Within individual cutting planes, AcD incidence was significantly larger in the hippocampal CA1 and CA3 areas compared to the subiculum. Interestingly, whereas central CA1 in the dorsal hippocampus shows the lowest incidence of AcD cells, their number reaches the highest levels in the medial and retains high levels in the ventral hippocampus. Here, they comprise around half of all pyramidal cells. We also found that AcD prevalence decreases from superficial to deep cell layers in ventral CA1 areas. We verified the different prevalence of AcD morphology between ventral and dorsal CA1 in the human brain, using 3 human samples aged 56 to 76 years old (**Figure 3C**). Cell morphology was visualized using antibodies against NeuN for the soma and βIV-spectrin for the AIS. Dorsal and ventral sections of the mouse hippocampus correspond to the posterior and anterior portions of the human hippocampus, respectively. Across all samples, we found the number of pyramidal cells with AcD morphology to be higher in the ventral/anterior sections, reproducing the pattern of the mouse hippocampus.

**Figure 3:**
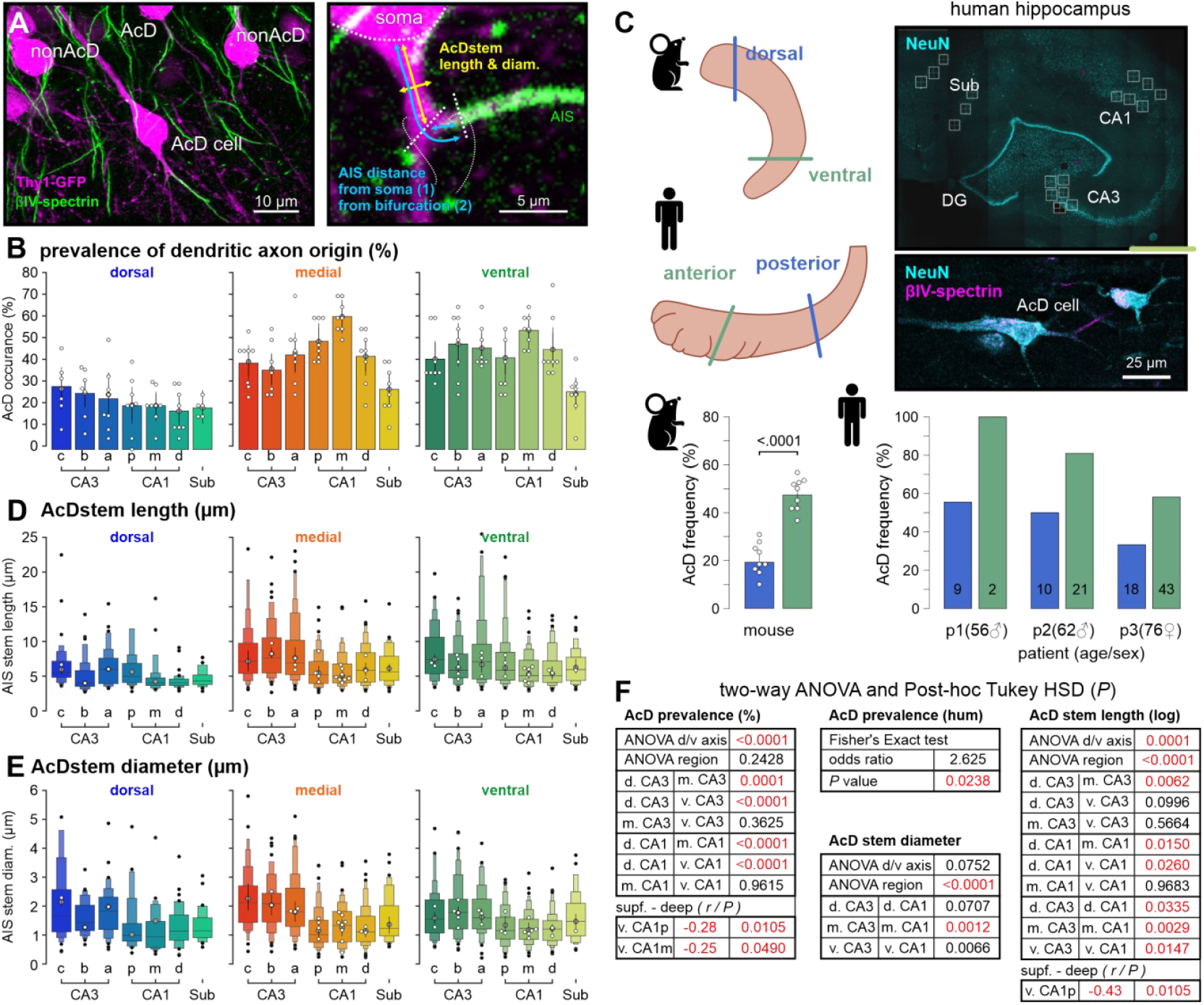
Dendritic axon origins are commonplace in the hippocampal formation. **(A)** Confocal image of pyramidal neurons with dendritic axon origins (AcD cells). If the segment separating the soma from the AIS-containing axon (AcD stem) was longer than its diameter (AcD stem diameter), the cell was classified as an AcD cell. **(B)** Bar plots illustrating the mean percentage of AcD cells among all cells in the respective region. Markers as in Figure 2A. **(C)** In human tissue, CA1 pyramidal cells frequently show AcD morphology with similar dorsal to ventral distribution as observed in mice. The top left panels illustrate orientation and homolog sections of the human and mouse hippocampus. Top right panels show confocal images of hippocampal formation from a human sample with neurons visualized by staining against NeuN and βIV-spectrin. Bottom panels show ratio of AcD to nonAcD cells in ventral/anterior and dorsal/posterior cutting sections of 8 mice (left) and 3 humans (right) of different ages. **(D & E)** Extended box plots of AcD stem lengths and mean diameters between different subregions and cutting planes of the hippocampal formation. For details on plot design see Figure 2A. **(F)** Statistical comparison between hippocampal areas and sections across the dorsal to ventral axes of the hippocampus. P values are derived from two-way ANOVA tests, following Tukey post-hoc analysis.

The functional relevance of AcD morphology is likely dependent on the longitudinal electrical resistance that separates the soma from the AIS (Fekete et al., 2021, Thome et al., 2014). We therefore measured the length and thickness of the AcD stem dendrite (**Figure 3D**), which separates the axon-carrying branches from the soma. Similar to the general propensity for AcD morphology, the median lengths of stem dendrites were shorter in the dorsal compared to the medial and ventral portions of the hippocampus across CA1 and CA3, while CA3 featured generally longer AcD stem dendrites compared to CA1 and subiculum. AcD stem diameters are similar along the dorsal-ventral axis but show considerable differences between hippocampal regions (**Figure 3E**). CA3 showed larger diameters compared to CA1 and in medial and ventral slices, also compared to the subiculum. AcD stem diameters in the subiculum were positioned between CA3 and CA1, though were only significantly different from both in the medial cutting plane. Superficial pyramidal cells tended to have longer AcD stems in medial and ventral sections, with highest discrepancies within ventral CA1 (*r*-value: −0.43 in proximal CA1 of ventral slices). A minor subpopulation had their AIS arising from the apical dendrite (<1%, data not shown).

### Morphological profiles

Hippocampal sub-regions exhibited individual profiles of proximal morphologies, in which cells belonging to major hippocampal regions were similar in shape (**Figure 4A**). Measures of AIS length, distance, and somatic/dendritic size were generally smaller in the dorsal hippocampus and larger in the intermediate cutting plane across all subregions, pointing to a systematic effect modulating the morphology of all principal neurons across the hippocampal longitudinal axis.

**Figure 4:**
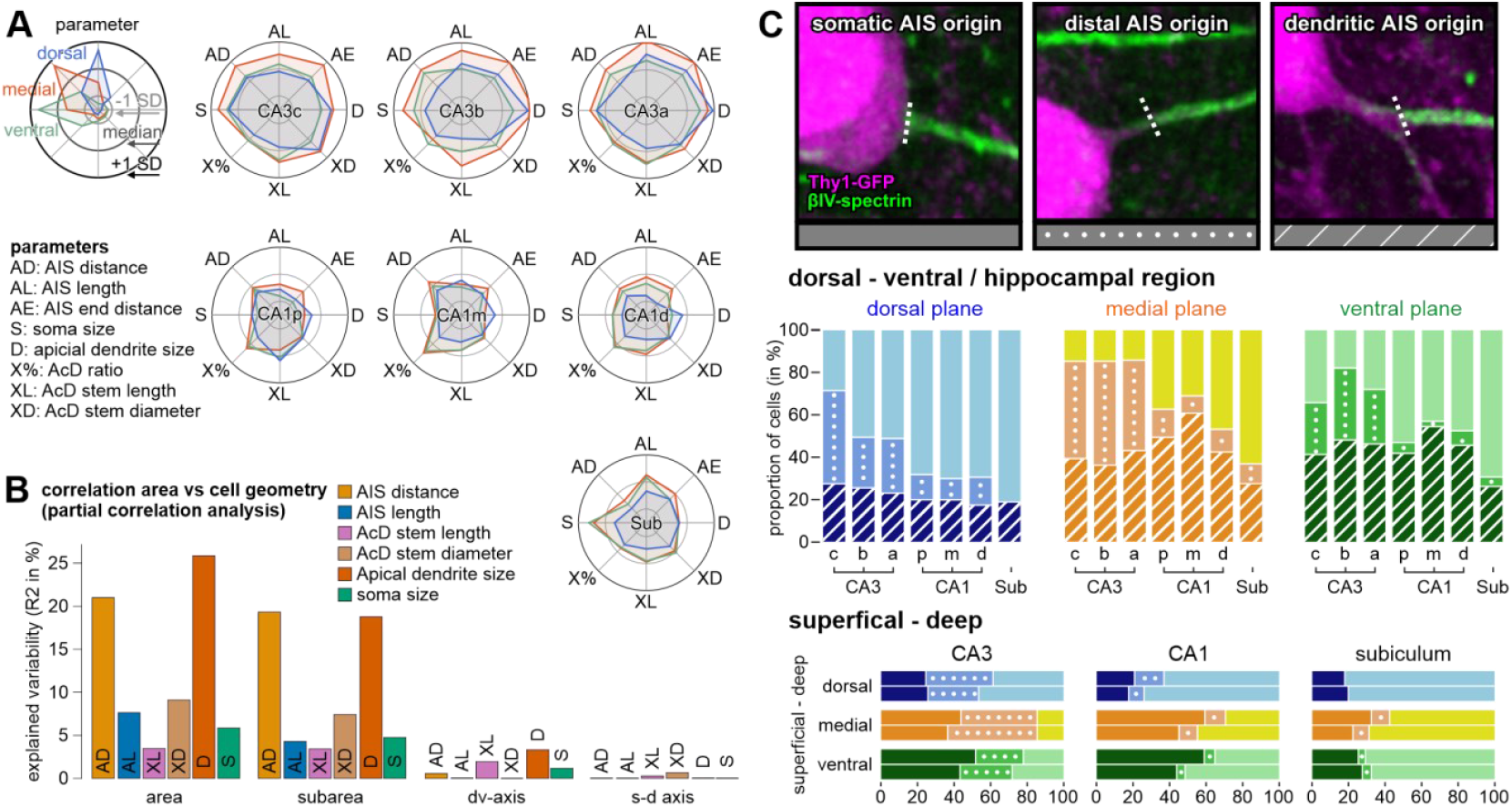
Cell morphology profiles across hippocampal subregions. **(A)** Radar chart plots depicting normalized relative medium values of morphological parameters. Circles show the median of all cells across all regions (middle circle), +/-standard deviation (outer/inner circle). Plots are color coded according to dorsal-ventral axes. **(B)** Partial correlation analysis visualizes how much of the observed variation in cell geometry could be predicted from the location of a given cell. Hippocampal areas and subareas have the most profound impact. **(C)** Classification of cell geometries in cells with somatic AIS origin, distal AIS origin (> 5µm), and dendritic AIS origin (AcD stem longer than thick). Bars show ratios of all animals and hippocampi pooled. Cells with AcD geometry were most abundant in medial and ventral hippocampal areas and showed generally lower numbers in the parahippocampal subiculum.

We used non-parametric pairwise partial correlation analysis to investigate the impact of cell location on proximal cell morphology (**Figure 4B)**. The hippocampal region had the biggest influence on cell anatomy, followed by the dorsal-ventral axis. The depth inside the cell layer had no general effect on any morphological parameter.

Pyramidal cells can be divided according to their axon origin into 3 different classes: neurons with somatic axon origin, distal axon origin (> 5 µm from the soma), and dendritic axon origin (Hofflin et al., 2017). These three morphologies were shown to have a significant impact on how neurons integrate dendritic input and generate action potential output. We thus investigated how these subgroups are distributed across the hippocampal formation **(Figure 4C)**. We found that the proportion of the three morphologies shows a gradual change along with the hippocampal subgroups and dorsal to ventral planes. As already described in Figure 3B, AcD cells are common across all regions, but their numbers are lower in the dorsal plane and peak in the medial and ventral regions. Distant axon origins that are not accompanied by AcD morphology were rare in most hippocampal subregions apart from CA3, independent of the respective amount of AcD morphology. Along the superficial-deep axis in dorsal slices, we observed a higher prevalence of cells with distant AIS in the upper layer of the pyramidal cell band. In contrast, their numbers remained consistent in the medial and ventral hippocampus, regardless of the increased occurrence of AcD morphology in superficial layers (see Figure 3F for comparison).

### Interdependencies of proximal cell geometries

It has previously been shown that in some regions, morphological parameters correlate in a compensatory way, leading to overall similar excitability (Hamada et al., 2016). We thus analyzed the interaction of different proximal cell geometries and confirmed established and expected interactions throughout all cell classes, like the positive correlation of cell size with AIS length **(Figure 5A)**. As expected, AcD stem length correlated strongly with the distance of the AIS, accounting for ∼78% of the measured variation (*P*< 0.000, *r* = 0.80, partial correlation controlled for cell location in hippocampal formation; **Figure 5B**). We now asked which parameters and interdependencies dominate AIS geometry across the hippocampal formation.

**Figure 5:**
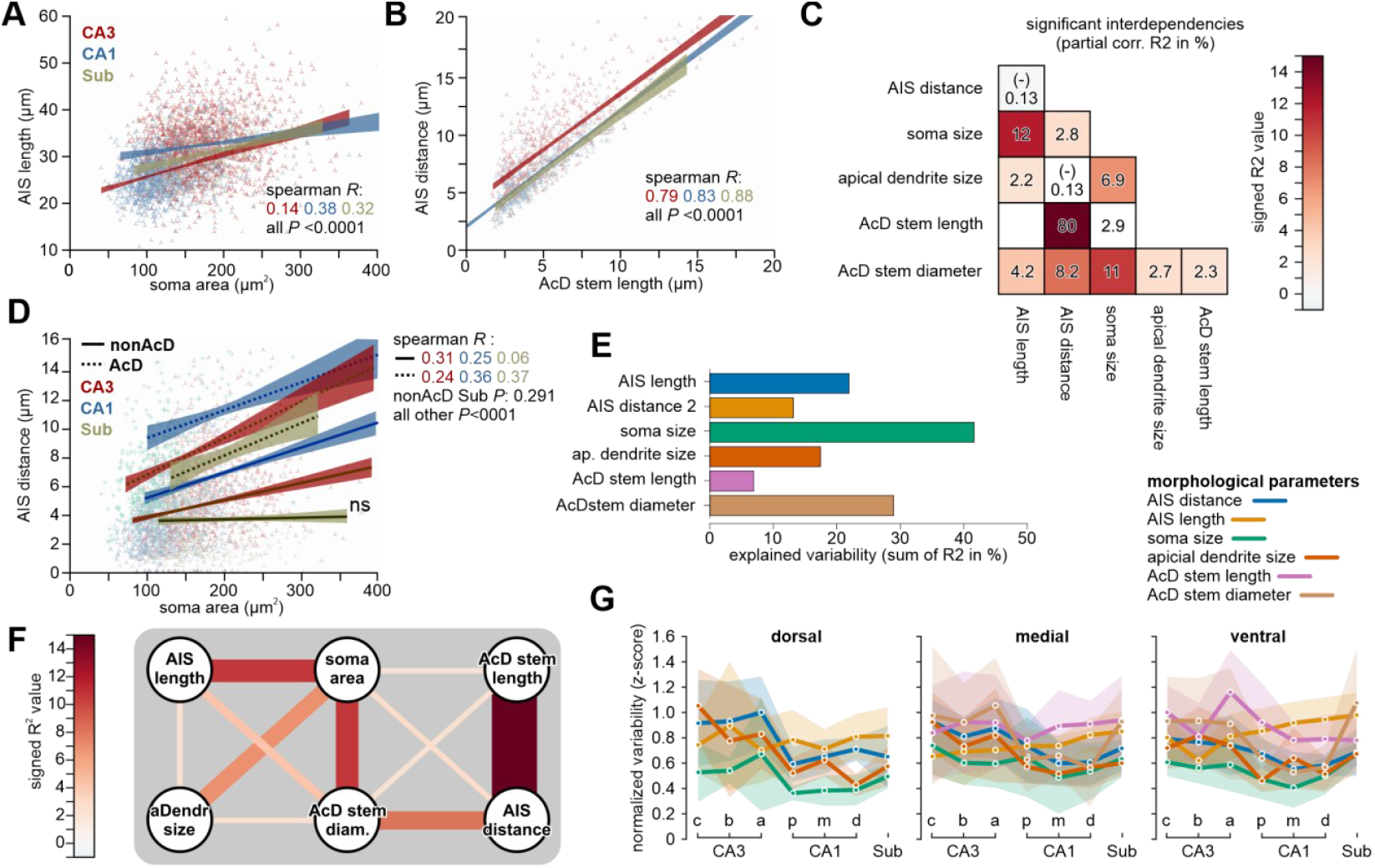
Interdependencies between parameters of cell geometry. **(A)** Scatter plot and linear regression analysis for significant correlations of AIS length with soma size in CA3, CA1, and subiculum. Shaded areas depict 95% confidence intervals. **(B)** Scatter plot and linear regression analysis for significant correlations of AIS stem length with AIS distance in CA3, CA1, and subiculum. **(C)** Partial correlation analysis of proximal cell geometry reveals several significant relationships between soma size and AIS geometry. The diameter of the AcD stem but not its length is tightly connected to the size of the soma. The length of the AcD stem shows remarkably little correlation with the other parameters of cell geometry (R2 values show percentage of predicted variability; partial correlation analysis from all CA1, CA3, and subicular cells; if not part of the analysis cell location was controlled for as covariates; 3326 cells from 12 animals). **(D)** Scatter plot and linear regression analysis for significant correlations of AIS distance with soma size in CA3, CA1, and subiculum, split between AcD and nonAcD cells. **(E)** The sum R2 for each parameter of cell geometry illustrates how well the variability of each parameter can be predicted from the others (partial correlation analysis). **(F)** Visualization of the strength of correlations with the color code given in in C. **(G)** Line plot of the variability of each morphological parameter across the hippocampal formation. Variability is computed as the SD of normalized (Z-scored) values for direct comparison across parameters. Confidence intervals reflect the standard deviation across hippocampi.

Using partial correlation analysis (including all measured parameters and cell positions) we distilled a general scheme of interdependencies subtracting the effect of anatomical confounding factors (including position in dorsal-ventral axis and cell type) (**Figure 5C**). The diameter of the AcD stem was more strongly correlated with other morphological parameters than the AcD stem length (apart from the inherently correlated AIS distance). It showed a strong interdependency with the size of the soma and was itself rather weakly correlated to the AcD stem length. Soma size correlated positively with the length of the AcD stem dendrite. However, soma size generally correlated with AIS distance which, in the case of AcD cells, is tightly correlated with AcD stem length. When we compared linear regression models of AIS distance and soma area in AcD cells vs. nonAcD cells, we found similar correlation coefficients for both groups in CA1 and CA3. Though in AcD cells the AIS were generally located at more distant positions (**Figure 5D**).

We proceeded with partial correlations to estimate how much variability in a given parameter can be predicted from the data we obtained about the other parameters. Because of the dominant intrinsic connection of AIS distance and AcD stem length in AcD cells, we included an alternative definition of AIS distance (AIS-distance-2, **Figure 3A**) in this analysis which describes the distance from the somatodendritic compartment (including the AcD stem) to the AIS. We found that around 45% of the variability of cell size can be predicted solely from the other measured parameters, with decreasing levels of predictability for AcD stem diameters and AIS lengths.

Interestingly, the length of AcD stem dendrites was most independent from the other parameters, indicating that variability in this parameter is only weakly counteracted by proximal cell morphology. This indicates that homeostatic compensation of AcD morphology would be less directed to the length of the AcD stem dendrite but rather its diameter, apart from the general and more dominant correlations associated with distal AIS origins. Note, that these statistical comparisons are blind to the actual physiological impact that individual parameters might contribute to the function of a given neuron.

We also analyzed the variability of morphological parameters across different locations in hippocampal planes **(Figure 5G)**. The dorsal axis shows a distinct pattern of variability, with the CA3 region displaying the highest variability across most parameters. This suggests significant structural diversity in this region. There is a noticeable decrease in variability from CA3 to CA1 regions. Along the medial axis, variability is more evenly distributed across the hippocampal regions, with a slight increase observed in the subregions. In the ventral axis, variability trends are more dispersed, with no single region showing markedly higher variability. Parameters of AcD morphology were excluded in the dorsal hippocampus due to their low incidence.

### Computer Predictions

We used our dataset to predict the variability in signal integration properties that may arise specifically from the proximal cell morphology. We were particularly interested in whether features that may increase neuronal excitability, such as AIS length and distal positions, were compensated by those that reduce excitability, such as larger capacitive sinks through the soma and apical dendrite as shown for other regions (Hamada et al., 2016).

Using a published computer model of a CA1 pyramidal cell, (Hines, 1984, Hines et al., 2009) we explored how the natural variability of proximal cell geometries might translate into a range of different neuronal excitabilities, disregarding other properties such as ion channel densities which were kept constant throughout our experiments. Based on the morphologies of all fully traced pyramidal cells, we constructed individual computer simulations of neurons with synaptic inputs at two basal dendrites (**Figure 6A**). We injected stochastic inputs into a simulated nonAcD branch and AcD branch (for AcD neurons) and determined the current threshold at which the cell fired an action potential. All cells fired action potentials within a realistic range of input currents (100-300 pA; **Figure 6B**) and voltage thresholds (−58 to −50 mV). Though medians of neuronal excitability were predicted to be different between different brain regions and dorsal to ventral axes, their distributions were strongly overlapping. We then evaluated the contribution of each geometric parameter to calculated input efficiencies in all CA1 pyramidal neurons (**Figure 6C**). While parameters of membrane size such as apical dendrite diameter and AcD stem diameter (but not soma size) had the strongest predictive power for input efficiency, the voltage threshold, measured at the soma, showed the highest correlation with AIS length. It has been established that neurons with and without AcD morphology do not differ in their gross neuronal excitability, but in how they integrate inputs from the AcD and nonAcD branches (Hamada et al., 2016, Hodapp et al., 2022, Thome et al., 2014). Our model cell was equipped with two basal dendritic branches of which one could be modified to feature the measured cell geometries. Injecting current in either branch of nonAcD cells resulted in equal firing thresholds for both branches. Dendritic branches of nonAcD cells with distal AIS origins featured higher current thresholds, but with a generally similar range of input efficiencies. In the case of AcD cells, both branches diverged in their ability to trigger action potentials, which broadened the range of input efficiencies without changing the size or ion channel composition of each dendrite. Note that a proximal section of the second nonAcD branch was always modified in length and diameter to keep symmetry between branches intact (see red halves in **Figure 6D**).

**Figure 6:**
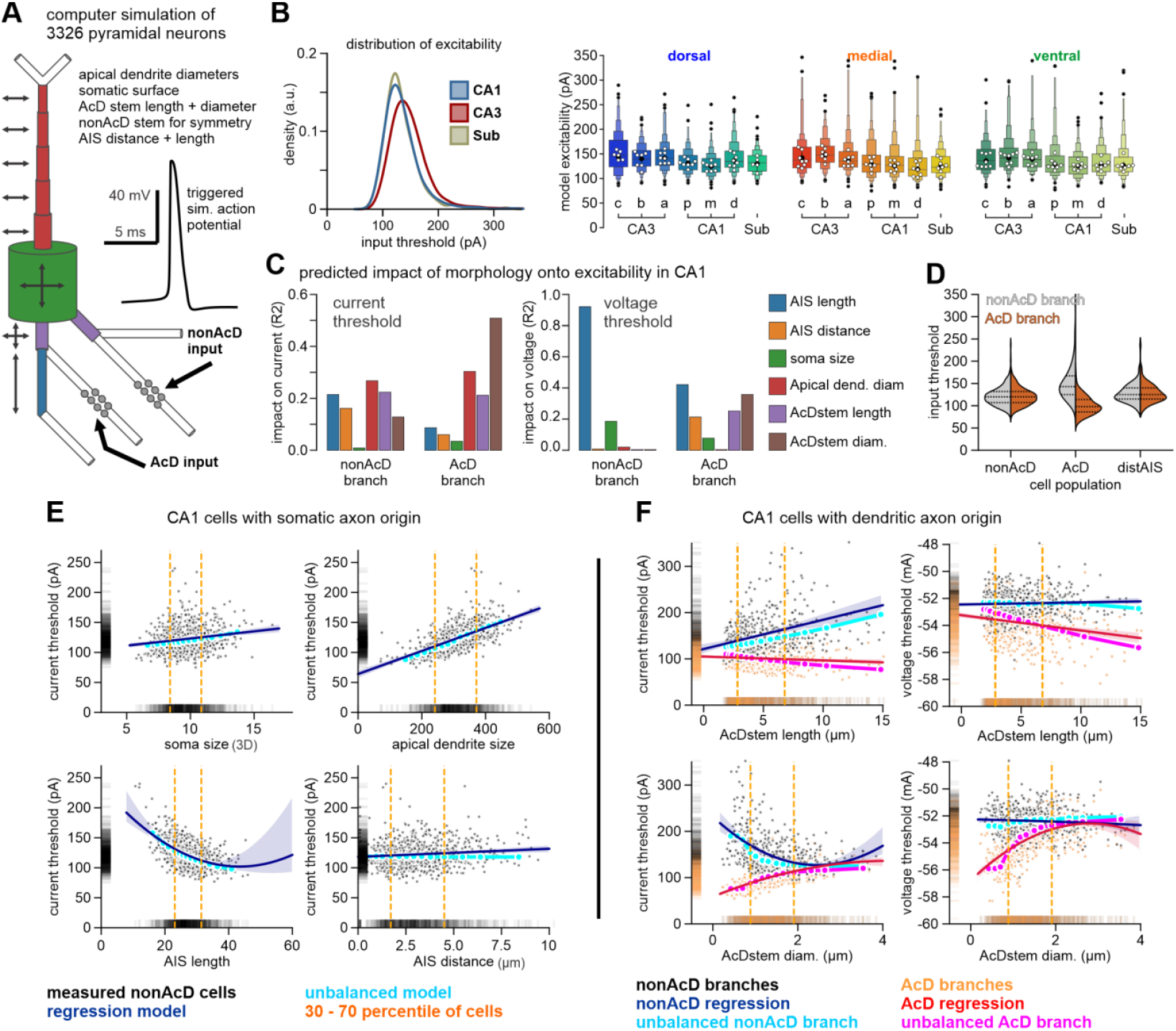
Computer simulation of neuronal excitability using measured cell geometries. **(A)** Cell compartments of a simulated neuron were modified according to manual measurements. Synaptic currents were injected either at the nonAcD or AcD type dendritic branch until the cell generated an action potential. For nonAcD cells, the AcD stem length was set to zero. **(B)** Mapping the predicted excitability for all measured cell geometries produced a wide range of current thresholds across all hippocampal subregions. **(C)** Using geometries of all CA1 pyramidal cells, the predicted current thresholds showed a large dependence on apical dendrite and AcD stem diameters, especially for the AcD branch (left panel). Voltage thresholds on the other hand were most tightly connected to AIS length though to a lower degree in AcD branches (middle panel). Bars are derived from partial correlation analysis. **(D)** Violin plots show the distribution of current thresholds between AcD and nonAcD branches of cell categories defined in Figure 4C. AcD morphology generated a considerable shift in the input efficiencies between the AcD and nonAcD branches of cells with dendritic axon origin. **(E)** Excitability of individual nonAcD cells ordered in relation to parameters of cell geometry (grey) compared to an idealized standard cell with static parameters (cyan, median values for all parameters of cell geometry). Linear regression of the dataset (dark blue) vs standard cell (cyan) shows a strong overlap, illustrating that there is only limited systematic homeostatic compensation by the other morphological parameters that could normalize input efficiencies. **(F)** The plot of current thresholds (left panels) and voltage thresholds (right panels) vs AcD stem lengths (top) and diameters (bottoms) illustrates the divergence of input efficiencies with increasing AcD morphology (longer and thinner AcD stem) between the AcD branch (orange dots) and nonAcD branches (grey dots). A standard cell featuring median values for all cell geometry but the one varied along the x-axis is plotted in cyan (nonAcD branch) and magenta (AcD branch). Regression analysis of the dataset shows a strong overlap with the unbalanced model cell, suggesting that there is limited systematic homeostatic compensation through other parameters of cell geometry. Second-order polynomial regression was used for AIS length in (E), as well as AcD stem diameter in (F).

While our model predicts a large variability in input efficiencies driven by proximal cell geometry, we could so far not establish whether individual geometries act mainly as homeostatic modulators, driving the cells back to a defined band of allowed excitability or as agents of diversification that boost variability in our simulation. To tackle this question, we defined a stereotypic CA1 pyramidal cell in which soma size, apical dendrite diameters, as well as AIS length, and position were kept at the medium value found in our dataset. We then modified only one of those parameters in the range of the 30% to 70% percentile of our dataset (**Figure 6E**, light blue dots). This experiment illustrates how our computer model predicts the impact of individual geometric cell parameters in general when all other parameters remain constant. We then compared the predicted current thresholds of model cells with the full cell geometries (**Figure 6E**, black dots). Applying simple regression models revealed that input efficiencies of cells with real geometries showed the same trajectory with e.g. AIS length as the stereotypic CA1 cell. Other correlations of cell geometries also followed the standard cell tightly, which lacks any potential homeostatic compensation. This was also true in AcD neurons for which we injected currents into the AcD and nonAcD branches of each neuron (**Figure 6F**).

We thus propose that the variability of AIS geometry has a dual impact on neuronal excitability. The interplay between AIS distance and AIS length on the one hand may compensate for somatic size and other anatomical and molecular mechanisms (see Figure 5). On the other hand, AIS geometry also increases neuronal diversity by enlarging the variability of neuronal excitability. The second case seems to be dominant in our model prediction although it is likely that other internal factors (e.g., ion channel composition, synaptic organization) as well as external factors (volume, position, and composition of excitatory and inhibitory drives) play a major role in setting the stage for the establishment and modulation of proximal cell geometries.

## DISCUSSION

### Summary

Understanding the structural variability of the AIS in hippocampal neurons is key for unraveling the mechanisms that underlie neuronal excitability and action potential generation. Our study provides a comprehensive analysis of AIS geometry across different regions of the murine hippocampus, focusing on how variations in AIS length, position, and other proximal cell geometries contribute to the functional diversity of neurons. By mapping these features across the dorsal-ventral, superficial-deep, and proximal-distal axes, we verified and uncovered significant regional differences that suggest a nuanced role of AIS geometry in shaping the unique functional characteristics of hippocampal neurons. The analysis provided in this publication does by no means exploit the full depth of the collected dataset, especially regarding patterns specific to neuronal subpopulations as well as CA2 and hilus. The authors invite all readers to use the provided dataset for their investigation.

### Limitations of the study

While our study provides significant insights into the structural diversity of AIS geometry and its potential functional implications, several limitations must be acknowledged. Due to the depth of our anatomical segmentation, we were limited in the number of animals and we do not have dorsal and ventral sections of the same animal due to our cutting procedure. Another limitation is the reliance on the Thy1-GFP mouse line for visualizing neuronal morphology. While this approach allowed for detailed structural analysis, it is selective for certain neuronal populations and may not represent the full diversity of AIS morphologies present in the hippocampus. CA2 neurons, for example, were Thy1-GFP negative. However, the distribution of AcD cells in this study confirms previous observations made with other methods (Hodapp et al., 2022). We furthermore compared the distributions of AIS lengths of Thy1-GFP-positive and negative cells of the same slices in CA1 and CA3 and found no differences.

Additionally, our study focused on the relationship between AIS geometry and proximal cell morphology, but did not directly assess the impact of other factors such as ion channel distribution, synaptic input patterns, or the influence of glial cells. All of these are known to affect neuronal excitability, but are difficult to access and quantify via immunofluorescence in fixed tissue. These limitations also transfer to our computer simulations. While they offer valuable predictions about the functional consequences of AIS geometry, they are based on simplified models that do not fully account for the complexity of real neuronal networks. For example, we used model parameters based on CA1 pyramidal cells of the dorsal hippocampus for all cell types and regions (**Figure 6B**), although these cells feature distinct molecular patterns and dendritic arborization. Complementing this dataset with functional recordings of neuronal excitability in undisturbed neurons could help validate and extend our findings.

### AIS diversity

Our analysis revealed substantial variability and overlap in AIS geometry across the murine hippocampal formation, with distinct profiles emerging in different subregions and along the primary anatomical axes. Dorsal cells generally exhibited the strongest difference in proximal cell anatomy compared to the other planes. These cells have smaller soma sizes, shorter and more distant initial segments, as well as fewer and less prominent axon-carrying dendrites across all hippocampal regions. AIS lengths and distances peaked in intermediate CA3, where the median AIS length was 35 µm and more than 80% of neurons had their axon origin >5 µm away from the soma. In the adjacent CA1 region, in contrast, initial segments were 10 µm shorter. Dendritic axon origins were most common in the intermediate hippocampus, where they are twice as common compared to dorsal CA1, a tendency that is observed in all subregions to some degree. These findings are consistent with previous studies that have reported regional differences in hippocampal neuron morphology but extend our understanding by providing a more granular view of AIS variability along multiple axes.

It is important to mention that while most AIS parameter shifts their average values along all three principal anatomical axes – [transverse (proximodistal), radial (deep-superficial), and longitudinal (dorsal-ventral)] - there remains a very strong overlap of the distributions (see Figure 2). The difference *within* each neuronal subpopulation is greater than the differences *between* them. One could say that the main hippocampal regions are more similar than dissimilar. This is important to keep in mind since many studies tend to visualize mean or median values that mask the real diversity of neuronal morphologies. Indeed, AIS heterogeneity may not only open up a large parameter space for baseline intrinsic excitability and modes of input integration. It may also allow for a differentially tuned susceptibility to AIS plasticity and subsequent modifications of neuronal excitability and firing precision (Hofflin et al., 2017, Rotterman et al., 2021).

Our findings suggest that neurons in different regions of the hippocampus have developed distinct AIS morphologies to meet the specific computational demands of their environment. For example, the longer and more distally positioned AIS in ventral CA3 neurons may compensate for their generally larger proximal somatodendritic surface, and facilitate quick responses to synaptic inputs. This increase in excitability may be crucial for the ventral hippocampus’ role in emotional processing and the rapid stress response, functions that are known to rely on robust and flexible neuronal firing patterns (Fanselow & Dong, 2010; Moser & Moser, 1998). Future studies using *in vivo* imaging techniques could provide a more comprehensive understanding of how AIS geometry correlates with neuronal activity under specific network conditions.

### Neuronal homeostasis

Apart from regional specificity, we found that region-specific differences in AIS length and distance are strongly associated with the size of the soma and apical dendrites. In the case of dendritic axon origins, the diameter of the AcD stem dendrite similarly correlated with AIS length. In general, the initial segment length and distance of the AIS correlate strongly with soma size, indicating a homeostatic balance that ensures a common regime of neuronal excitabilities within each subregion. Such morphological relationships are well in line with AIS plasticity rules described in previous studies, where neurons underwent shifts in AIS lengths and positions in response to changes in neuronal activity (Grubb and Burrone, 2010, Jamann et al., 2021, Kuba, 2012). However, we found no such relationship regarding the propensity and strength of AcD morphology. Cells with dendritic axon origins featured AIS distances up to 10 µm away from the soma, which was not accompanied by additional morphological characteristics apart from the ones that modulate or compensate for distant AIS locations in nonAcD neurons. A study in neocortical pyramidal cells showed that action potential waveforms in neocortical pyramidal cells with dendritic axon origins are normalized by a narrower apical dendrite (Hamada et al., 2016). In our dataset, we found no correlation between apical dendrite diameters and AIS distance from the soma in AcD or nonAcD neurons. Thus, the reported relationship may be specific to rat cortical infragranular pyramidal neurons and not present within the hippocampal formation of mice.

The impact of the length and position of the AIS on neuronal excitability was mostly described in the context of plasticity or computational studies (Goethals and Brette, 2020), but also (Rotterman et al., 2021). If AIS plasticity modulates excitability, homeostatic compensation should be limited. In our dataset, interactions of proximal cell geometries explain ∼7% (AcD stem length) up to ∼41% (soma size) of the found variation in cell morphology (**Figure 5E**). However, we cannot conclude to which degree these correlations attribute to homeostasis or increase the variability of neurons. We thus built a NEURON simulation environment in which we studied the theoretical impact of individual morphological parameters on neuronal excitability and how they are balanced within each cell. Other modulatory parameters, such as particular ion channels densities were kept static to evaluate the impact of cell morphology alone. While we received a wide distribution of input efficiencies, reminiscent of data from physiological recordings (Thome et al., 2024), solely based on our measured proximal cell geometry, they showed no signs of homeostatic balancing **(Figure 6E and 6F)**. Of course, if anatomy was always balanced to yield the same mean excitability, all pyramidal neurons regardless of shape or region would show the same firing thresholds. Electrophysiological data from in vitro and in vivo recordings demonstrate that this is far from the truth and likely incompatible with physiological network states (Pena and Rotstein, 2022). Indeed, cortical neurons exhibit large variability in excitability within each region, though the average of individual distributions may be region-specific.

### AcD neurons

Our study confirms and expands upon previous findings that AcD neurons exhibit no major differences regarding their general morphology or excitability, but reveal distinct properties during specialized tasks (Hamada et al., 2016, Hodapp et al., 2022, Stevens et al., 2024, Thome et al., 2014). We were nevertheless surprised that AcD morphology caused no evident deviation from the general pattern of correlations of cell geometry seen in nonAcD neurons. While AcD cells are found in the neocortex and also show a unique mode of input integration properties in dorsal CA1 (Hodapp et al., 2022), the highest density of AcD morphology is documented in the intermediate and ventral hippocampus. We can only speculate about the functional implications of this pattern.

CA1 pyramidal cells in the ventral hippocampus feature higher neuronal excitabilities in patch clamp recordings, but at the same time have fewer dendritic branches and total dendritic surface area (Milior et al., 2016, Dougherty et al., 2012, Malik et al., 2016). These conditions may facilitate the formation of a privileged dendritic branch that integrates inputs of particular salience. This region is also associated with emotional aspects of memory processing which may involve stronger interhemispheric connectivity in turn associated with AcD cells in ventral CA1. Future studies will be necessary to confirm the actual input-output relation of neurons with different AIS morphologies under in vivo conditions. Furthermore, it will be crucial to verify whether AcD cells participate differently in network activities that might underlie the behavioral and functional segregation from the dorsal to the ventral end of the hippocampus. Moreover, given the translational potential of our human data, future research should also investigate the implications of AIS variability in the context of neurological disorders and aging. Variations in AIS structure have been linked to conditions such as epilepsy, schizophrenia, and autism spectrum disorder (Buffington and Rasband, 2011, Ferreira et al., 2008, Fujitani et al., 2021, Garrido, 2023, Hsu et al., 2014, Huang and Rasband, 2018, Usui et al., 2022, Wimmer et al., 2010). Therefore, confirming our mouse data in human tissue as well as understanding how AIS geometry changes in human disease states and how these changes affect neuronal function could lead to novel therapeutic strategies targeting the AIS or related cellular structures.

## MATERIAL AND METHODS

### Animals and sample preparation

Animal housing was performed at the Interfaculty Biomedical Facility (IBF) at Heidelberg University, approved by the local animal welfare officer, and all procedures were performed in strict accordance with the guidelines of the European Community Council and the state government of Baden-Wurttemberg. We used male and female individuals of the Thy1-GFP mouse line (M) aged 3-4 months. Animals were anesthetized with CO_2_ and decapitated after the loss of the righting reflex. The brain was quickly removed and transferred into ice-cold (1–2°C) artificial cerebrospinal fluid (ACSF) containing the following (in mM): 124 NaCl, 3 KCl, 1.8 MgSO_4_, 1.6 CaCl_2_, 10 glucose, 1.25 NaH_2_PO_4_, saturated with 95% O_2_/5% CO_2_ and calibrated to pH 7.4. The cerebellum and the first third of the frontal brain were removed, and 350 μm horizontal slices were cut using a vibrating blade microtome (Leica VT1000S, Leica Biosystems). Coronal slices were made for dorsal hippocampal sections. Sections were approximately 750 µm apart.

### Immunofluorescence and confocal imaging

Slices were fixed in 2% paraformaldehyde (PFA; Sigma-Aldrich) diluted in 0.1 M phosphate buffer, pH 7.4, at room temperature for 90 min. After fixation, slices were stored in phosphate-buffered saline (PBS; in mM, 137 NaCl, 2.7 KCl, 8 NaH_2_PO_4_, pH 7.2) at 4°C. Axon initial segments were visualized using the primary antibody rabbit-anti-βIV-spectrin (1:1,000; self-made, corresponding to amino acids 2,237–2,256 of human βIV-spectrin; (Gutzmann et al., 2014)) combined with secondary antibody Alexa 647 antirabbit 1:1,000 (Molecular Probes). The staining procedure was performed as follows: slices were washed, blocked for 2 h in blocking solution (5% normal goat serum from Vector Laboratories, 0.5% Triton X-100 from Merck in PBS; washed again; and incubated in primary antibody solution (0.2% Triton X-100 and 1% NGS in PBS) overnight at room temperature. Slices were again washed and incubated for 2 h in the secondary antibody solution. All washing steps were done three times for 15 min in PBS and incubation steps were done on an orbital shaker. After the final washing step, slices were mounted on microscope slides using Fluoroshield (Abcam, Danaher Corporation) with DAPI to visualize the cell band. Slices were imaged with a Nikon A1R confocal microscope. We used objectives with magnification factors 10 (Nikon Plan Apo λ 10× NA 0.45) for overviews and 60x (Nikon N Apo 60× NA 1.4 λ oil immersion) confocal stacks of 0.5 μm z resolution to capture cell geometries. Human samples were imaged using a Leica Stellaris 5 confocal microscope with 10x and 63x objectives. Images were analyzed in 3D using ImageJ (Wayne Rasband, NIH, open source) and the simple neurite tracer package (Arshadi et al., 2021).

### Anatomical segmentation

Slices for morphological analysis were selected after staining visually based on anatomical characteristics. Ventral slices are the first sections that feature all hippocampal regions and have a semi-circular shape of dentate gyrus and pyramidal cell band. Medial slices were defined as the last slices before the pyramidal cell band was distorted by dorsal curvature, featuring a V-shaped dentate gyrus and pronounced subiculum. Samples for the dorsal hippocampus were selected for having a horizontal dentate gyrus and CA3c base. Anatomical subregions were defined according to cell morphology, location, and orientation as follows: hilus: disorganized cells between dentate gyrus blade; CA3c: pyramidal cells in the first 30 μm of the CA3 cell band; CA3b: cells in the curvature of the band; CA3a: cells at the end of CA3 but outside CA2; CA23: The lack of GFP in CA2 in our mouse line was confirmed by other labs (personal communication). We verified the CA2 position using the marker PCP4 which consistently labeled neurons around the border of stratum lucidum (Stevens et al., 2024). CA1 was split into proximal (start), central, and distal before transition to subiculum. Confocal images were taken in the center of each defined subregion without overlap. We aimed to capture 20 fully analyzed cell geometries per subregion. If cell numbers were not matched with the initial 4 animals, additional animals were used. This applied especially for CA3 areas in the dorsal planes. Superficial and deep cells were defined post hoc by their geometric distance towards the stratum radiatum border. Superficial layers were defined as ∼2 neurons wide and considering the total thickness of the cell band (30-50 μm; **Figure 1)**.

### Cell Classification and Tracing

The soma size was defined as convex curvature around the Thy1-GFP label. Cell size was captured as an ellipse defined by the two longest orthogonal diameters. The thickness of apical dendrites was measured by 5 consecutive width measurements starting from the base of the somatic circumference with 5 μm spacing. AIS start was measured from the somatic curvature until a strong increase in the βIV-spectrin fluorescence. The end was defined as the last point of consecutive labeling. We tried automated approaches using thresholds in fluorescence intensities to define the start and end of initial segments, but these were error-prone in our hippocampal dataset. In cases where the axon and AIS was connected to a somatic protrusion that also gave rise to a dendrite, we measured the length and diameter of this segment (AcD stem). Cells were only classified as AcD cells during post hoc data processing when the potential AcD stem was at least 2 μm long and longer than its mean diameter.

### Human samples

Samples of hippocampal tissue were provided by the Department of Neurosurgery, Kepler University Hospital Linz in strict accordance with the approved ethical protocol. Fixation and staining were performed as described above, but using fish gelatine instead of NGS as a blocking agent.

### Computer Simulations

Computer simulations were carried out using the NEURON simulation environment through its python module (Hines et al., 2009). Time step duration was set to 25 µs (corresponding to 40 kHz). The soma gave rise to a single apical dendrite and two basal dendrites. The apical dendrite was divided into three parts with different diameters and ion channel composition and terminated into two distal dendritic branches mimicking an apical tuft. One of the basal dendrites was connected to the AIS (AcD branch), while the other (non-AcD branch) carried another dendritic structure to achieve electrotonic symmetry with the AcD branch. The origin of the third dendrite was shifted along the non AcD branch in parallel to the axon distance to maintain the electrotonic symmetry between AcD and non AcD. The AIS was constructed out of two parts with the activation threshold of Na^+^ channels in the distal part shifted by −5 mV to simulate experimental findings (Hu et al., 2009b). The passive electrical properties Rm, Cm, and Ra were set to 25,000 Ωcm2, 1 µF/cm2, and 200 Ωcm, respectively. Segmentation around the dendritic injection sites was made with high resolution (0.5 µm) to keep high-frequency terms of the input signal. Active conductances were modified from previously reported values (Thome et al., 2014, Cutsuridis et al., 2010, Hu et al., 2009a). The conductances of the individual segments were similar to (Hodapp et al., 2022).

### Statistical Methods

Cell morphologies were collected from up to 20 individual cells per subregion per hippocampus and pooled for analysis. The resulting distributions exhibited either symmetrical bell-shaped or log-normal characteristics (e.g., AcD distance, AcD stem length). Log-normal distributions were logarithmically transformed before applying statistical tests such as ANOVA; otherwise, nonparametric tests were employed. Statistical significance was set at *P* < 0.05 for all tests, with significant p-values denoted in figures by asterisks (< 0.05*, < 0.01**, < 0.001***). All analyses were conducted using the Python module Pingouin (Vallat, 2018). One statistical limitation warrants attention: cells and hippocampi were treated as independent values in all statistical analyses, though this is not entirely accurate. We aimed to sample 20 fully characterized pyramidal cells per location within each anatomical subregion. In instances of low GFP expression, additional animals were included to achieve approximately equal cell counts across subregions. Notably, medial and ventral samples were derived from the same animals, while dorsal slices were prepared from separate individuals. To assess the influence of sex, hemisphere, and animal on cell geometry parameters, we performed 3-way ANOVAs, finding no significant differences.

Consequently, values from each hippocampal location with more than nine fully measured cells are represented as individual data points in Figures 2 and 3. While we consider this approach suitable for descriptive purposes and controlled for potential confounds using partial correlation tests (Figure 5), it may affect the precision of statistical tests used in Figure 2 (comparison of hippocampal planes and main areas). For further reference, animal identifiers, sex, and hemisphere information are included in the dataset. The R2 value was used to assess the strength of the correlations between morphological parameters and anatomical location. Specifically, it quantifies the proportion of variance in one variable that can be predicted by another.

To assess and compare the variability of six morphological parameters across different hippocampal regions, we first applied a log transformation to AIS distance and AcD stem length to address their non-normal distribution. Subsequently, we normalized all parameters globally by calculating Z-scores across the entire dataset. Variability was then computed for each hippocampus as the standard deviation of the Z-scored values within each hippocampal region and anatomical axis. The variability data was visualized using either line plots with confidence intervals based on the standard deviation of variability across individuals or letter-value plots. These extended box plots (Letter-value plots, boxenplots in pythons seaborn package) show distribution of AIS length, AIS distances, somatic area, and mean apical dendrite diameter between area CA1, CA3, and subiculum. White dots depict median values of the region for each hippocampus. Grey dots and lines depict mean value of all hippocami pooled with 95% confidence interval. The black dots depict outliers as defined by data points falling beyond the extended ranges determined by the letter-value plot method. Specifically, these ranges are dynamically calculated based on the interquartile range (IQR) at progressively smaller proportions of the data distribution, following the ‘k_depth’ parameter set to ‘trustworthy.’ Outliers represent values that deviate significantly from the main distribution.

## Author contributions

C.T. conceived and designed the research project. C.T. and N.A.S. conducted the experimental research, with N.A.S. tracing all cell morphologies. M.B. developed the computer simulations. R.L. provided the human samples, which were stained by J.M. and traced by M.A. Data analysis was performed by C.T., who also drafted the initial version of the manuscript. All authors reviewed and contributed to the final manuscript. This work was supported by the German Research Foundation (DFG BO 3512/2-1 to M.B., DFG EN 1240/2-1 to M.E., and the Walter Benjamin Programme PN 458054460 to C.T.).

## Supporting information

dataset and python code

## Acknowledgements

The authors thank Tina Sackmann and Katja Lankisch for their support with animal handling, Nadine Zuber for her contribution with staining and solution preparation; and the Nikon Imaging Center Heidelberg and the Core Facility Imaging of JKU Linz for providing the microscopes. A special thank is extended to Prof. Dr. Andreas Draguhn, for providing the lab, resources, and guidance to support this project.

## SUPPLEMENTAL MATERIAL

**Supplemental Figure S1:**
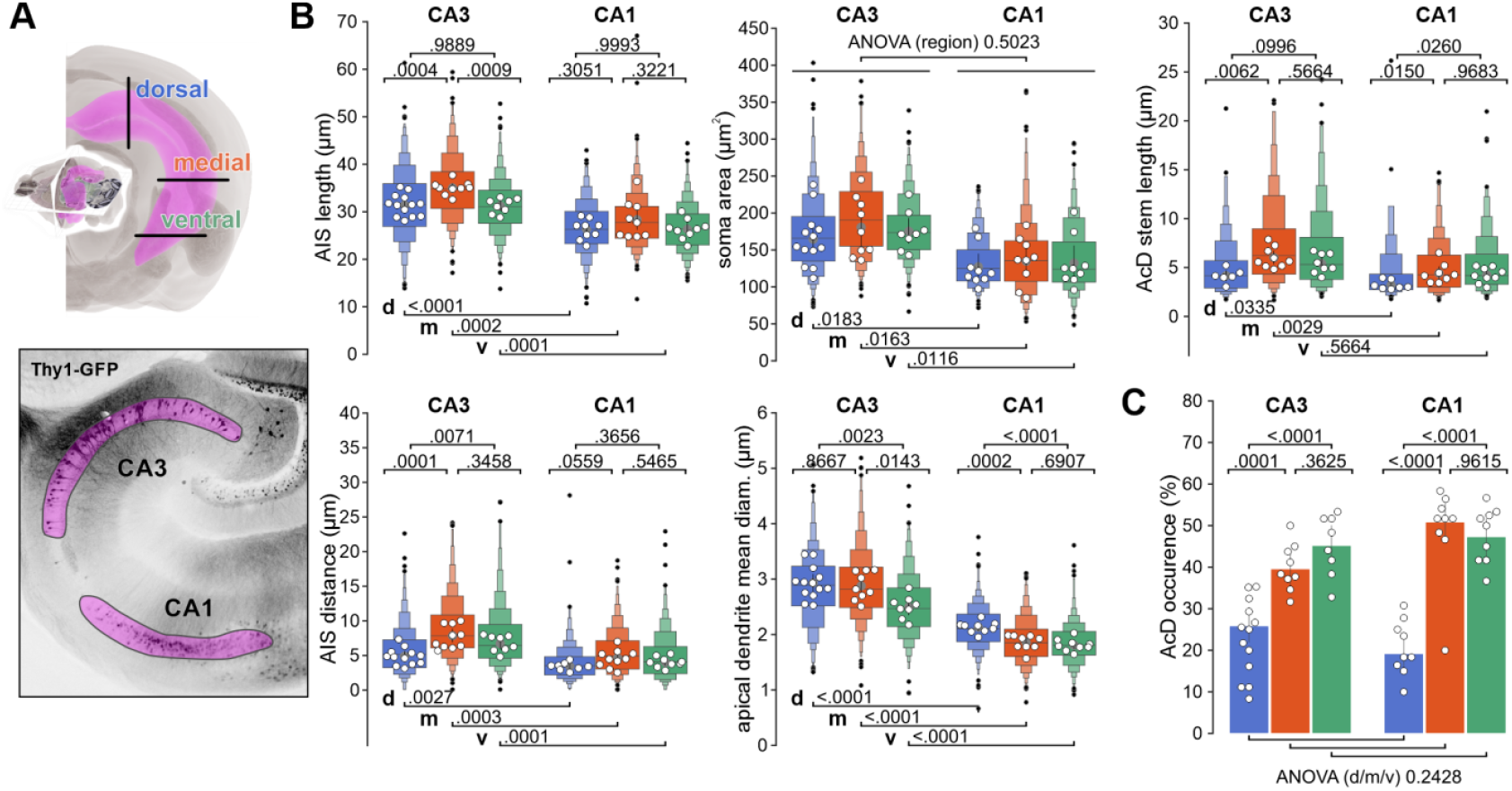
Comparison of proximal cell parameters between hippocampal regions in different cutting planes along the dorsal-ventral axis. **(A)** Scheme of murine hippocampus (Allen Brain Explorer) illustrating color code of hippocampal sections used in this study (top panel). Bottom panel: Confocal image of Thy1-GFP signal in ventral slice. CA1 and CA3 areas are marked by magenta areas. **(B)** Extended box plots (Letter-value plots, boxenplots in pythons seaborn package) show distribution of AIS length, AIS distances, soma area, and mean apical dendrite diameter between area CA3 and CA1. White dots depict median values of the region for each hippocampus. Grey dots and lines depict mean value of all hippocami pooled with 95% confidence interval. The black dots depict outliers as defined by data points falling beyond the extended ranges determined by the letter-value plot method. Specifically, these ranges are dynamically calculated based on the interquartile range (IQR) at progressively smaller proportions of the data distribution, following the ‘k_depth’ parameter set to ‘trustworthy.’ Outliers represent values that deviate significantly from the main distribution. Statistical differences were determined by performing an ANOVA to test for differences along the dorsal-to-ventral axis or between hippocampal regions (CA3 and CA3). In cases where significantly different distributions were identified, a Tukey post-hoc test was conducted to determine pairwise differences. *P*-values are indicated above the respective boxes in the figure. **(C)** Barplots depicting the mean percentage of dendritic axon origins across hippocampal regions and across different cutting planes depicted in (A). Markers and statistics are as described in (B).

**Supplemental Figure S2:**
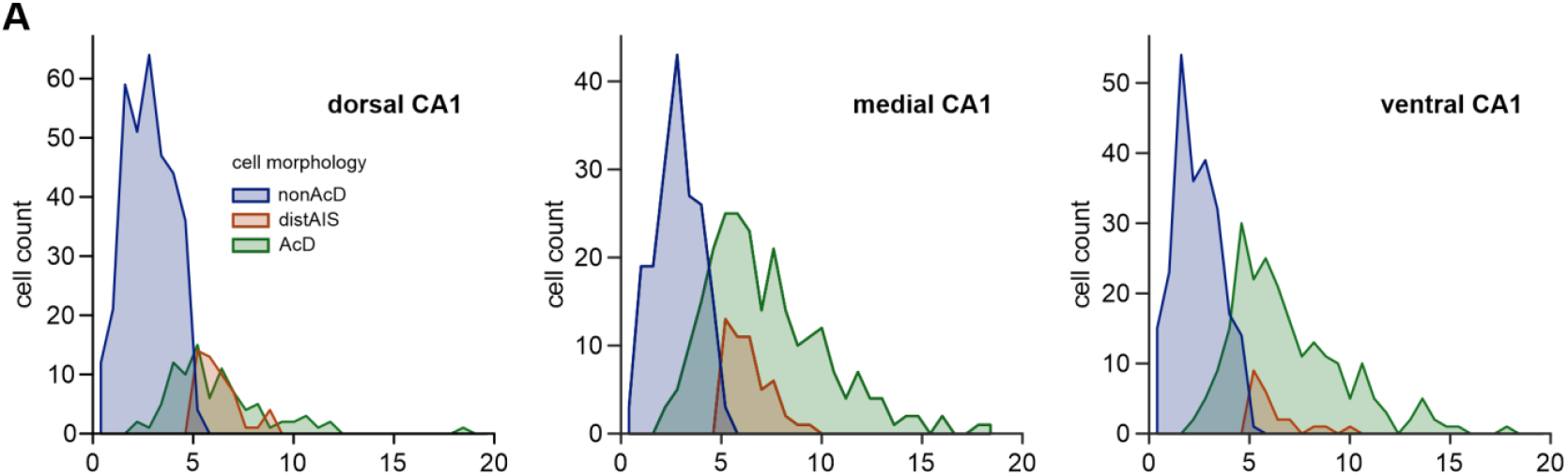
Distribution of AIS distances in CA1 pyramidal neurons. **(A)** Histograms show distribution of AIS distances in CA1 pyramidal cells separated by AIS origin. Compare with Hodapp et al., 2022.

**Supplemental Figure S3:**
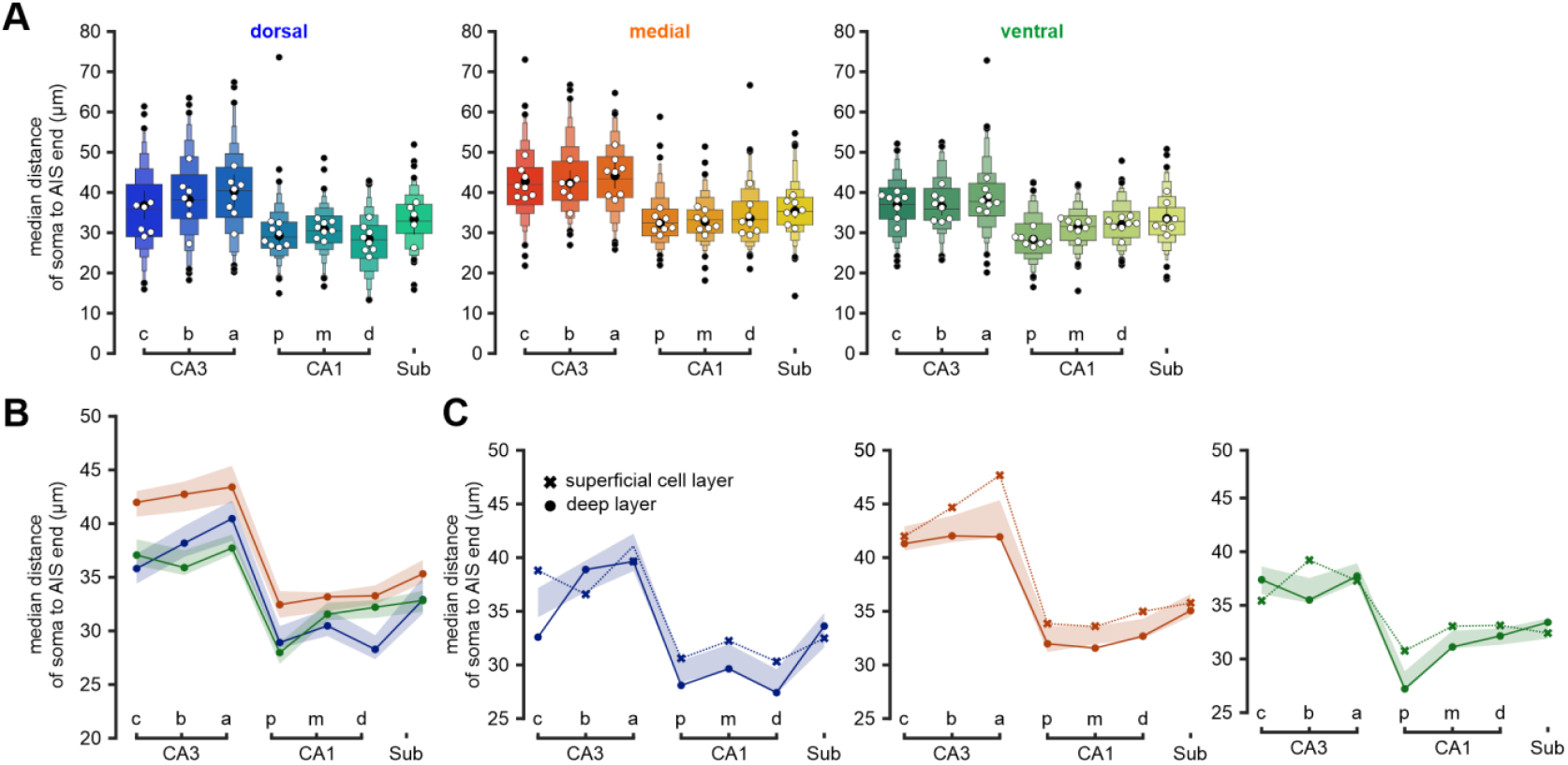
Comparison of the distances between soma and the end of the AIS label βIV-spectrin between hippocampal subregions in different cutting planes along the dorsal-ventral axis. **(A)** Extended box plots designed as described in Figure S2A. **(B)** Line plots show the median distance of the end of the AIS fluorescence signal as in A for better comparison between dorsal, medial, and ventral axes (left panel). **(C)** Line plots show the median distance of the end of the AIS fluorescence signal as in A and B but split according to dorsal-ventral axes and further divided by the cell position within the pyramidal cell band (x: superficial, o: deep).

