## Supplementary material for "Variability of axon initial segment geometry and its impact on hippocampal pyramidal cell function": dataset and python code: AISmorph_corr-matrices_plot.py

```
################## READ ME FIRST ######################
# the code needs the following python packages to run: pandas, seaborn, numpy, matplotlib, os
# the code uses the provided excel sheet 'AISmorph_masterlist.xlsx' which should be saved somewhere on your computer
# the code will plot 4 correlation matrices between all morphological parameter and create a JPEG file in the same folder as the source excel file
# the numbers inside the boxes depict signed R2 values of only statitistcally significant correlations
# the matrices are compiled from different selections of cells. 
# Top left: all cells; Top right: all cells but AcD cells; Bottom right: only AcD cells; Bottom left: only cells with AIS distances > 5um
# the code needs two inputs from the users:

# 1st: define the path were the 'AISmorph_masterlist.xlsx' is stored on your hard drive in the line 'excelpath' (line 12)
# 2nd: comment areas you want to be excluded from the analysis (place '#' sign in front of all other parameters or they might become overwritten)


###################    USER INPUT AREA  ######################

################ SET FILE PATH ##############################

excelpath = r"G:\Meine Ablage\Y_Projects\Y_AISdiversity_PYR_mouse\submission\bioarchive"

################ SET PARAMETER FOR X AXIS ##############################

################ EXCLUDE HIPPOCAMPAL REGIONS OR CELLTYPES ##############################

areas_to_filter = [
    ('hilus', 'hcarea2'),
    # ('CA3', 'hcarea2'),
    ('CA2', 'hcarea2'),
    # ('CA1', 'hcarea2'),
    # ('Sub', 'hcarea2'),
    
    # ('CA3c', 'hcarea'),
    # ('CA3b', 'hcarea'),
    # ('CA3a', 'hcarea'),
    # ('CA1p', 'hcarea'),
    # ('CA1m', 'hcarea'),
    # ('CA1d', 'hcarea'),
    
    # ('celltype', 'AcD'),
    # ('celltype', 'nonAcD'),
    # ('celltype', 'distAIS'),

]


#%% import packages and data
import pandas as pd
import seaborn as sns
import matplotlib.pyplot as plt
import os
import pingouin as pg
import numpy as np

#%% import masterfile

rawfile = os.path.join(excelpath,'AISmorph_masterlist.xlsx')
datasheet = pd.read_excel (rawfile,sheet_name='cellmorphologies') 

#%% define variables

# Initialize a mask for rows to exclude
exclude_mask = pd.Series(False, index=datasheet.index)

# Build the exclusion mask
for area_name, column_name in areas_to_filter:
    if column_name in datasheet.columns:  # Check if the column exists
        exclude_mask |= (datasheet[column_name] == area_name)
    else:
        print(f"Warning: Column '{column_name}' not found in the DataFrame. Skipping '{area_name}'.")

# Filter the data by excluding rows matching the conditions
filtered_data = datasheet[~exclude_mask]
filtered_data2 = filtered_data[['AISlength','AISdist', 'AISdist2',
                   'soma_area','area_Adend',
                   'AcDstem','AcDstem_diam',
                   'hcarea2_ind','dvaxis_ind','supfic_binary_ind',
                   'hcarea2','dvaxis','supfic_binary',  'celltype',
                   'AcDness',
                   'animal_ind']] 


#%%
fig1 = plt.figure(figsize=(10,8), constrained_layout=True)
gs0 = fig1.add_gridspec(2, 2)
ax1  = fig1.add_subplot(gs0[0,0]) # all cells
ax2  = fig1.add_subplot(gs0[0,1]) # nonAcD cells 
ax3  = fig1.add_subplot(gs0[1,0]) # AcD cells
ax4  = fig1.add_subplot(gs0[1,1]) # nonAcD cells with distant AIS
sns.despine()

areastitle = filtered_data.hcarea.unique()
title_text = "Analysis includes neurons of regions:\n" + ", ".join(areastitle)
fig1.suptitle(title_text, fontsize=14, weight='bold', ha='center')


#%% plot1 for all cells
data1 = filtered_data2
# Define your morphological variables and covariates
mophvar_list1 = ['AISlength','AISdist','AISdist2', 
                   'soma_area','area_Adend',
                    'AcDstem','AcDstem_diam'
                   ]
covar_list1 = ['hcarea2_ind','dvaxis_ind']

# Initialize an empty DataFrame to store adjusted R² values
r_squared_signed_matrix = pd.DataFrame(index=mophvar_list1, columns=mophvar_list1)

# Compute partial correlations for each pair of morphological variables
for var1 in mophvar_list1:
    for var2 in mophvar_list1:
        if var1 != var2:
            # Calculate partial correlation
            result = pg.partial_corr(data=data1, x=var1, y=var2, covar=covar_list1)
            p_value = result['p-val'].values[0]
            correlation = result['r'].values[0]
            
            # Only store the signed R² value if the p-value is significant
            if p_value < 0.05:
                signed_r_squared = (correlation ** 2) * np.sign(correlation)  # Include the sign
                r_squared_signed_matrix.loc[var1, var2] = signed_r_squared
            else:
                r_squared_signed_matrix.loc[var1, var2] = np.nan  # or 0 to avoid displaying it in the heatmap
        else:
            r_squared_signed_matrix.loc[var1, var2] = 1  # Diagonal should be 1 as R² of a variable with itself is 1

# Convert to float and round the values
r_squared_signed_matrix = r_squared_signed_matrix.astype(float).round(4) * 100

# Plot the heatmap of signed R² values with normalized colormap
mask = np.zeros_like(r_squared_signed_matrix)
mask[np.triu_indices_from(mask)] = True
sns.heatmap(r_squared_signed_matrix, 
            cmap='RdBu_r',  # Divergent colormap for signed values
            mask=mask, square=True, center=0, 
            vmin=-1, vmax=15,  # Normalize colormap from -15 to +15
            linewidths=.5, annot=True, ax=ax1)

ax1.set_title('all cells (signed R²)')
ax1.set_xticklabels(labels=mophvar_list1, rotation=20)
ax1.set_yticklabels(labels=mophvar_list1, rotation=20)


####################################

#%% Plot2 for nonAcD cells

data1 = filtered_data2
data1 = data1[data1.celltype != 'AcD']

mophvar_list1 = ['AISlength', 'AISdist',
                  'soma_area', 'area_Adend']
covar_list1 = ['hcarea2_ind', 'dvaxis_ind']

r_squared_matrix = pd.DataFrame(index=mophvar_list1, columns=mophvar_list1)

for var1 in mophvar_list1:
    for var2 in mophvar_list1:
        if var1 != var2:
            result = pg.partial_corr(data=data1, x=var1, y=var2, covar=covar_list1)
            p_value = result['p-val'].values[0]
            correlation = result['r'].values[0]
            
            if p_value < 0.05:
                signed_r_squared = (correlation ** 2) * np.sign(correlation)
                r_squared_matrix.loc[var1, var2] = signed_r_squared
            else:
                r_squared_matrix.loc[var1, var2] = np.nan
        else:
            r_squared_matrix.loc[var1, var2] = 1

r_squared_matrix = r_squared_matrix.astype(float).round(4) * 100

mask = np.zeros_like(r_squared_matrix)
mask[np.triu_indices_from(mask)] = True
sns.heatmap(r_squared_matrix, cmap='RdBu_r', mask=mask, square=True, 
            vmin=-1, vmax=15, center=0, linewidths=.5, annot=True, ax=ax2)

ax2.set_title('nonAcD cells (signed R²)')
ax2.set_xticklabels(labels=mophvar_list1, rotation=20)
ax2.set_yticklabels(labels=mophvar_list1, rotation=20)

#%% Plot3 for AcD cells only

data1 = filtered_data2
data1 = data1[data1.celltype == 'AcD']

mophvar_list1 = ['AISlength', 'AISdist','AISdist2', 
                  'soma_area', 'area_Adend',
                    'AcDstem','AcDstem_diam']
covar_list1 = ['hcarea2_ind', 'dvaxis_ind']

r_squared_matrix = pd.DataFrame(index=mophvar_list1, columns=mophvar_list1)

for var1 in mophvar_list1:
    for var2 in mophvar_list1:
        if var1 != var2:
            result = pg.partial_corr(data=data1, x=var1, y=var2, covar=covar_list1)
            p_value = result['p-val'].values[0]
            correlation = result['r'].values[0]
            
            if p_value < 0.05:
                signed_r_squared = (correlation ** 2) * np.sign(correlation)
                r_squared_matrix.loc[var1, var2] = signed_r_squared
            else:
                r_squared_matrix.loc[var1, var2] = np.nan
        else:
            r_squared_matrix.loc[var1, var2] = 1

r_squared_matrix = r_squared_matrix.astype(float).round(4) * 100

mask = np.zeros_like(r_squared_matrix)
mask[np.triu_indices_from(mask)] = True
sns.heatmap(r_squared_matrix, cmap='RdBu_r', mask=mask, square=True, 
            vmin=-1, vmax=15, center=0, linewidths=.5, annot=True, ax=ax3)

ax3.set_title('AcD cells (signed R²)')
ax3.set_xticklabels(labels=mophvar_list1, rotation=20)
ax3.set_yticklabels(labels=mophvar_list1, rotation=20)


#%% Plot4 for distal AIS only

data1 = filtered_data2
data1 = data1[data1.celltype == 'distAIS']

mophvar_list1 = ['AISlength', 'AISdist',
                  'soma_area', 'area_Adend']
covar_list1 = ['hcarea2_ind', 'dvaxis_ind']

r_squared_matrix = pd.DataFrame(index=mophvar_list1, columns=mophvar_list1)

for var1 in mophvar_list1:
    for var2 in mophvar_list1:
        if var1 != var2:
            result = pg.partial_corr(data=data1, x=var1, y=var2, covar=covar_list1)
            p_value = result['p-val'].values[0]
            correlation = result['r'].values[0]
            
            if p_value < 0.05:
                signed_r_squared = (correlation ** 2) * np.sign(correlation)
                r_squared_matrix.loc[var1, var2] = signed_r_squared
            else:
                r_squared_matrix.loc[var1, var2] = np.nan
        else:
            r_squared_matrix.loc[var1, var2] = 1

r_squared_matrix = r_squared_matrix.astype(float).round(4) * 100

mask = np.zeros_like(r_squared_matrix)
mask[np.triu_indices_from(mask)] = True
sns.heatmap(r_squared_matrix, cmap='RdBu_r', mask=mask, square=True, 
            vmin=-1, vmax=15, center=0, linewidths=.5, annot=True, ax=ax4)

ax4.set_title('nonAcD cells with distal AIS (signed R²)')
ax4.set_xticklabels(labels=mophvar_list1, rotation=20)
ax4.set_yticklabels(labels=mophvar_list1, rotation=20)


#%% export figure to output path
output_file = f"{excelpath}/plot correlation matrices.jpeg"
fig1.savefig(output_file, format='jpeg', dpi=300)
```
