## Supplementary material for "Variability of axon initial segment geometry and its impact on hippocampal pyramidal cell function": dataset and python code: AISmorph_correlations_plot.py

```
################## READ ME FIRST ######################
# the code needs the following python packages to run: pandas, seaborn, numpy, matplotlib, os
# the code uses the provided excel sheet 'AISmorph_masterlist.xlsx' which should be saved somewhere on your computer
# the code will plot a descriptive graph of a given morphological parameter and create a JPEG file in the same folder as the source excel file
# the code needs two inputs from the users:

# 1st: define the path were the 'AISmorph_masterlist.xlsx' is stored on your hard drive in the line 'excelpath' (line 12)
# 2nd: Uncomment the desired morphological parameter with associated limits for the Y axes by removing the '#'
    # dont forget to comment (place '#' sign in front of all other parameters or they might become overwritten)


###################    USER INPUT AREA  ######################

################ SET FILE PATH ##############################

excelpath = r"G:\Meine Ablage\Y_Projects\Y_AISdiversity_PYR_mouse\submission\bioarchive"

################ SET PARAMETER FOR X AXIS ##############################

x_parameter = ['AISlength', 'AIS length', (0, 60)]
# x_parameter = ['AISdist', 'AIS distance', (-8, 30)]
# x_parameter = ['logAISdist', 'log AIS distance', (-3, 4)]
# x_parameter = ['AcDstem', 'AcD stem length', (0, 25)]
# x_parameter = ['AcDstem_diam', 'AcD stem diameter', (0, 6)]
# x_parameter = ['soma_area', 'Soma area', (0, 400)]
# x_parameter = ['Adend_diam_mean', 'Apical dendr. diameter', (0, 5)]
# x_parameter = ['AcD_nes', 'AcD frequency', (0, 0.8)]  
# x_parameter = ['nAcD_ampa', 'Non-AcD excitability', (0, 350)]
# x_parameter = ['AcD_ampa', 'AcD excitability', (0, 350)]
# x_parameter = ['nAcD_Vthres', 'Non-AcD voltage threshold', (-60, -48)]
# x_parameter = ['AcD_Vthres', 'AcD voltage threshold', (-60, -48)]
# x_parameter = ['AISend', 'AIS end', (0, 80)]

################ SET PARAMETER FOR X AXIS ##############################

# y_parameter = ['AISlength', 'AIS length', (0, 60)]
# y_parameter = ['AISdist', 'AIS distance', (-8, 30)]
# y_parameter = ['logAISdist', 'log AIS distance', (-3, 4)]
# y_parameter = ['AcDstem', 'AcD stem length', (0, 25)]
# y_parameter = ['AcDstem_diam', 'AcD stem diameter', (0, 6)]
y_parameter = ['soma_area', 'Soma area', (0, 400)]
# y_parameter = ['Adend_diam_mean', 'Apical dendr. diameter', (0, 5)]
# y_parameter = ['AcD_nes', 'AcD frequency', (0, 0.8)]  
# y_parameter = ['nAcD_ampa', 'Non-AcD excitability', (0, 350)]
# y_parameter = ['AcD_ampa', 'AcD excitability', (0, 350)]
# y_parameter = ['nAcD_Vthres', 'Non-AcD voltage threshold', (-60, -48)]
# y_parameter = ['AcD_Vthres', 'AcD voltage threshold', (-60, -48)]
# y_parameter = ['AISend', 'AIS end', (0, 80)]

################ EXCLUDE HIPPOCAMPAL REGIONS OR CELLTYPES ##############################

areas_to_filter = [
    ('hilus', 'hcarea2'),
    # ('CA3', 'hcarea2'),
    ('CA2', 'hcarea2'),
    # ('CA1', 'hcarea2'),
    # ('Sub', 'hcarea2'),
    
    # ('CA3c', 'hcarea'),
    # ('CA3b', 'hcarea'),
    # ('CA3a', 'hcarea'),
    # ('CA1p', 'hcarea'),
    # ('CA1m', 'hcarea'),
    # ('CA1d', 'hcarea'),
    
    # ('celltype', 'AcD'),
    # ('celltype', 'nonAcD'),
    # ('celltype', 'distAIS'),

]

groups_to_compare = str(
    # 'hcarea2',
    'hcarea',
    # 'celltype',
    # 'supfic',
    )

#%% import packages and data
import pandas as pd
import seaborn as sns
import matplotlib.pyplot as plt
import os
from scipy.stats import spearmanr
#%% import masterfile

rawfile = os.path.join(excelpath,'AISmorph_masterlist.xlsx')
datasheet = pd.read_excel (rawfile,sheet_name='cellmorphologies') 


#%% define variables
orderlist = ["CA3c", "CA3b", "CA3a", "CA1p", "CA1m", "CA1d", "Sub"]
hcarea2_list = ["CA3", "CA1", "Sub"]
dvaxis_list = ["dorsal", "medial", "ventral"]
whitedots = sns.color_palette(['white', 'white', 'white'])

datasheet['AISdist2'] = datasheet.AISdist
datasheet.loc[datasheet.celltype == 'AcD', 'AISdist2'] = datasheet.AISdist - datasheet.AcDstem

# Initialize a mask for rows to exclude
exclude_mask = pd.Series(False, index=datasheet.index)

# Build the exclusion mask
for area_name, column_name in areas_to_filter:
    if column_name in datasheet.columns:  # Check if the column exists
        exclude_mask |= (datasheet[column_name] == area_name)
    else:
        print(f"Warning: Column '{column_name}' not found in the DataFrame. Skipping '{area_name}'.")

# Filter the data by excluding rows matching the conditions
filtered_data = datasheet[~exclude_mask]

# Compute Spearman correlation for each group and add annotations to the plot
def compute_and_annotate_spearman(filtered_data, x_param, y_param, group_column, ax):
    # Store results for exporting
    correlation_results = []

    # Iterate over unique groups
    unique_groups = filtered_data[group_column].unique()
    for group in unique_groups:
        group_data = filtered_data[filtered_data[group_column] == group]

        # Compute Spearman correlation
        r, p = spearmanr(group_data[x_param], group_data[y_param])

        # Round r and handle p-values
        r_rounded = round(r, 4)
        p_formatted = "<0.0001" if p < 0.0001 else round(p, 4)

        # Annotate the plot with the Spearman correlation in the lower-left corner
        ax.text(
            0.05, 0.05 + 0.05 * list(unique_groups).index(group),  # Adjust position for lower-left corner
            f"{group}: r={r_rounded}, p={p_formatted}",
            transform=ax.transAxes, fontsize=8, verticalalignment='bottom'
        )

        # Append results to the list
        correlation_results.append({
            "Group": group,
            "Spearman_r": r_rounded,
            "P-value": p if p >= 0.0001 else "<0.0001"
        })

    # Return results as a DataFrame
    return pd.DataFrame(correlation_results)

#%% compute correlations


#%% figure head
fig1 = plt.figure(figsize=(5, 5), constrained_layout=True)
gs0 = fig1.add_gridspec(1, 1)  # Adjusted grid to include ax6
ax1 = fig1.add_subplot(gs0[0, 0])
sns.despine()
#%% PLOT1  soma vs AIS length 

# Scatterplot with hue for groups
sns.scatterplot(
    x=x_parameter[0], y=y_parameter[0], hue=groups_to_compare, 
    palette='dark', alpha=0.3, marker='o', s=15, ax=ax1,
    data=filtered_data
)

# Dynamically add regplots for each group in the hue
unique_groups = filtered_data[groups_to_compare].unique()  # Get unique group names
for group in unique_groups:
    sns.regplot(
        x=x_parameter[0], y=y_parameter[0],
        color=sns.color_palette('dark')[list(unique_groups).index(group)],  # Get the color for the group
        truncate=True, robust=False, marker='', scatter=False, ax=ax1,
        data=filtered_data[filtered_data[groups_to_compare] == group]
    )

# Set labels and limits
ax1.set_xlabel(x_parameter[1])
ax1.set_ylabel(y_parameter[1])
ax1.set_xlim(x_parameter[2])
ax1.set_ylim(y_parameter[2])
# Compute Spearman correlations and annotate
correlation_df = compute_and_annotate_spearman(filtered_data, x_parameter[0], y_parameter[0], groups_to_compare, ax1)


#%% export figure to output path
output_file = f"{excelpath}/plot {x_parameter[1]} vs. {y_parameter[1]}.jpeg"
fig1.savefig(output_file, format='jpeg', dpi=300)
```
