## Supplementary material for "Variability of axon initial segment geometry and its impact on hippocampal pyramidal cell function": dataset and python code: AISmorph_morphology_plot.py

```
################## READ ME FIRST ######################
# the code needs the following python packages to run: pandas, seaborn, numpy, matplotlib, os
# the code uses the provided excel sheet 'AISmorph_masterlist.xlsx' which should be saved somewhere on your computer
# the code will plot a descriptive graph of a given morphological parameter and create a JPEG file in the same folder as the source excel file
# the code needs two inputs from the users:

# 1st: define the path were the 'AISmorph_masterlist.xlsx' is stored on your hard drive in the line 'excelpath' (line 12)
# 2nd: Uncomment the desired morphological parameter with associated limits for the Y axes by removing the '#'
    # dont forget to comment (place '#' sign in front of all other parameters or they might become overwritten)


###################    USER INPUT AREA  ######################

excelpath = r"G:\Meine Ablage\Y_Projects\Y_AISdiversity_PYR_mouse\submission\bioarchive"

parameter = ['AISlength', 'AIS length', (0, 60)]
# parameter = ['AISdist', 'AIS distance', (-8, 30)]
# parameter = ['logAISdist', 'log AIS distance', (-3, 4)]
# parameter = ['AcDstem', 'AcD stem length', (0, 25)]
# parameter = ['AcDstem_diam', 'AcD stem diameter', (0, 6)]
# parameter = ['soma_area', 'Soma area', (0, 400)]
# parameter = ['Adend_diam_mean', 'Apical dendr. diameter', (0, 5)]
# parameter = ['AcD_nes', 'AcD frequency', (0, 0.8)]  
# parameter = ['nAcD_ampa', 'Non-AcD excitability', (0, 350)]
# parameter = ['AcD_ampa', 'AcD excitability', (0, 350)]
# parameter = ['nAcD_Vthres', 'Non-AcD voltage threshold', (-60, -48)]
# parameter = ['AcD_Vthres', 'AcD voltage threshold', (-60, -48)]
# parameter = ['AISend', 'AIS end', (0, 80)]


#%% import packages and data
import pandas as pd
import seaborn as sns
import matplotlib.pyplot as plt
import numpy as np
import os

#%% import masterfile

rawfile = os.path.join(excelpath,'AISmorph_masterlist.xlsx')
datasheet = pd.read_excel (rawfile,sheet_name='cellmorphologies') 
AISmorph_statistics = pd.read_excel (rawfile,sheet_name='descriptive_stats') 

#%% define variables
# Assign the selected parameter
mphvar, mphvar2, yaxeslim = parameter
data2 = AISmorph_statistics
orderlist = ["hilus","CA3c","CA3b","CA3a","CA2","CA1p","CA1m","CA1d","Sub"]
datasheet = datasheet.sort_values(by='hcarea', key=lambda x: pd.Categorical(x, categories=orderlist, ordered=True))

if mphvar == 'AIS length':
    data1 = datasheet
else:
    data1 = datasheet[datasheet.hcarea !='CA2'][datasheet.hcarea !='hilus']
hcarea_list = ["CA3c","CA3b","CA3a","CA1p","CA1m","CA1d","Sub"]
hcarea2_list = ["hilus","CA3","CA2","CA1","Sub"]
dvaxis_list = ["dorsal","medial","ventral"]
oderlist = hcarea_list
xvalue = "hcarea"
whitedots  = sns.color_palette(['white','white','white'])

statavg ='median'
estimator = np.median
if mphvar == 'AcD_nes': 
    statavg = 'mean'
    estimator = np.mean
    data1 = data1[data1.hcarea2 != 'CA2']

#%%  figure head

fig1 = plt.figure(figsize=(12,8),constrained_layout=True)
gs0 = fig1.add_gridspec(2, 3)
ax1  = fig1.add_subplot(gs0[0,0])
ax2  = fig1.add_subplot(gs0[0,1])
ax3  = fig1.add_subplot(gs0[0,2])
ax4  = fig1.add_subplot(gs0[1,0])
ax5  = fig1.add_subplot(gs0[1,1])
ax6  = fig1.add_subplot(gs0[1,2])
sns.despine()

#%% figures individual plots
####################
plane = "dorsal"
if mphvar == 'AcD_nes':
    sns.barplot(x=xvalue, y=mphvar,palette="winter",
                   order=oderlist,  ax=ax1,\
                  data=data1[data1.dvaxis==plane])
else:
    sns.boxenplot(x=xvalue, y=mphvar,palette="winter", k_depth='trustworthy',
                   order=oderlist, showfliers=True, ax=ax1,\
                       flier_kws=dict(facecolor="k", linewidth=.5),
                   data=data1[data1.dvaxis==plane])
sns.pointplot(x=xvalue, y=statavg, color='.5',\
              order=oderlist, join=False, ax=ax1, 
              data=data2[data2.dvaxis==plane][data2.variable==mphvar])
sns.swarmplot(x=xvalue, y=statavg,ax=ax1, 
              order=oderlist, color='white',edgecolor='black',  linewidth=1,
              data=data2[data2.dvaxis==plane][data2.number>9][data2.variable==mphvar])
ax1.set(ylim=yaxeslim)
ax1.title.set_text(plane+ ' Hippocampus')
ax1.set_xlabel('')
ax1.set_ylabel(mphvar2)
#ax1.set_yscale('log')


########################
plane = "medial"
if mphvar == 'AcD_nes':
    sns.barplot(x=xvalue, y=mphvar,palette="autumn", 
                  order=oderlist,  ax=ax2,\
                  data=data1[data1.dvaxis==plane])
else:
    sns.boxenplot(x=xvalue, y=mphvar,palette="autumn",k_depth='trustworthy',
              order=oderlist, showfliers=True, ax=ax2,\
                       flier_kws=dict(facecolor="k", linewidth=.5),
              data=data1[data1.dvaxis==plane])
sns.pointplot(x=xvalue, y=statavg, color='.5',\
              order=oderlist, join=False, ax=ax2,\
              data=data2[data2.dvaxis==plane][data2.variable==mphvar])
sns.swarmplot(x=xvalue, y=statavg,ax=ax2, 
              order=oderlist, color='white',edgecolor='black',  linewidth=1,
              data=data2[data2.dvaxis==plane][data2.number>9][data2.variable==mphvar])
ax2.set(ylim=yaxeslim)
ax2.title.set_text(plane+ ' Hippocampus')
ax2.set_xlabel('')
#ax2.set_yscale('log')
ax2.set_ylabel(mphvar2)
########################
plane = "ventral"
if mphvar == 'AcD_nes':
    sns.barplot(x=xvalue, y=mphvar,palette="summer",
                  order=oderlist,  ax=ax3,\
                  data=data1[data1.dvaxis==plane])
else:
    sns.boxenplot(x=xvalue, y=mphvar,palette="summer",k_depth='trustworthy',
              order=oderlist, showfliers=True, ax=ax3,\
                       flier_kws=dict(facecolor="k", linewidth=.5),
              data=data1[data1.dvaxis==plane])
sns.pointplot(x=xvalue, y=statavg, color='.5',
              order=oderlist, join=False, ax=ax3,\
              data=data2[data2.dvaxis==plane][data2.variable==mphvar])
sns.swarmplot(x=xvalue, y=statavg,ax=ax3, 
              order=oderlist, color='white', edgecolor='black',  linewidth=1,
              data=data2[data2.dvaxis==plane][data2.number>9][data2.variable==mphvar])
ax3.set(ylim=yaxeslim)
ax3.title.set_text(plane+ ' Hippocampus')
ax3.set_ylabel(mphvar)
ax3.set_xlabel('')
#ax3.set_yscale('log')
ax3.set_ylabel(mphvar2)
###################

#%% line plots
# Map the colors for each category in 'dvaxis' to ensure consistency
line_palette = sns.color_palette("dark", len(dvaxis_list))
dvaxis_colors = dict(zip(dvaxis_list, line_palette))

# Iterate through the dvaxis categories and create separate subplots
for ax, dvaxis, label in zip([ax4, ax5, ax6], dvaxis_list, ["dorsal", "medial", "ventral"]):
    subset = data1[data1['dvaxis'] == dvaxis]  # Filter data for the current dvaxis
    sns.lineplot(
        x="hcarea",
        y=mphvar,
        hue="dvaxis",
        palette=dvaxis_colors,
        estimator=estimator,
        sort=False,
        linewidth=0.0,
        ax=ax,
        legend=False,
        data=subset
    )
    sns.lineplot(
        x="hcarea",
        y=mphvar,
        hue="dvaxis",
        style="supfic_binary",
        palette=dvaxis_colors,
        markers=True,
        errorbar=None,
        estimator=estimator,
        # linewidth=0.5,
        ax=ax,
        legend=False,
        data=subset
    )
    ax.set_title(label + ' Hippocampus')
    ax.set_ylabel(statavg + " " + mphvar2)
    ax.set_xlabel("hcarea")
    ax.set_ylim(yaxeslim)
    if mphvar in ['AIS length', 'AISend']: 
        yaxeslim = (20, 50) 
        ax4.set(ylim=yaxeslim)

#%% export figure to output path
output_file = f"{excelpath}/plot {mphvar2}.jpeg"
fig1.savefig(output_file, format='jpeg', dpi=300)
```
